# Coordinated control of genome-nuclear lamina interactions by Topoisomerase 2B and Lamin B receptor

**DOI:** 10.1101/2024.10.01.616012

**Authors:** Stefano G Manzo, Tom van Schaik, Marcel de Haas, Jeremie Breda, Mikhail Magnitov, Elzo de Wit, Anna G Manjon, Rene H Medema, Adam J Buckle, Catherine Naughton, Nick Gilbert, Bas van Steensel

## Abstract

Lamina-associated domains (LADs) are megabase-sized genomic regions anchored to the nuclear lamina (NL). Factors controlling the interactions of the genome with the NL have largely remained elusive. Here, we identified DNA topoisomerase 2 beta (TOP2B) as a regulator of these interactions. TOP2B binds predominantly to inter-LAD (iLAD) chromatin and its depletion results in a partial loss of genomic partitioning between LADs and iLADs, suggesting that its activity might protect specific iLADs from interacting with the NL. TOP2B depletion affects LAD interactions with lamin B receptor (LBR) more than with lamins. LBR depletion phenocopies the effects of TOP2B depletion, despite the different positioning of the two proteins in the genome. This suggests a complementary mechanism for organising the genome at the NL. Indeed, co-depletion of TOP2B and LBR causes partial LAD/iLAD inversion, reflecting changes typical of oncogene-induced senescence. We propose that a coordinated axis controlled by TOP2B in iLADs and LBR in LADs maintains the partitioning of the genome between the NL and the nuclear interior.

**Highlights:**

- LADs and iLADs differ in supercoiling state
- TOP2B controls genome partitioning between nuclear lamina and nuclear interior
- TOP2B depletion preferentially affects genome interactions with LBR
- Similar impact of TOP2B depletion and LBR depletion on genome-NL interactions
- Co-depletion of TOP2B and LBR recapitulates LAD reshaping typical of oncogene-induced senescence.

## INTRODUCTION

The typical nuclear organisation of metazoan cells requires an extensive portion of the genome (up to 40%) to interact with the nuclear lamina (NL) (5–7). Such interactions occur within extended regions of heterochromatin defined as lamina-associated domains (LADs). LADs have been extensively studied over the years, revealing that these structures have low gene density, harbor lowly expressed genes and heterochromatic marks such as H3K9 and H3K27 methylation, and exhibit transcriptional repressive capacity (5 Briand, 2020, 32241294,8-10). However, despite a good characterisation of LAD chromatin features, the key regulators of chromatin association with the NL remain poorly described.

The heterochromatic state of the LAD and its direct regulators have been proposed to play a role in genome-NL association. For example, H3K9me2 is a mark that is enriched at the nuclear periphery and is conserved across species (11). Tethering G9a, the main H3K9me2 writer, to a specific target sequence causes its association with the NL (12). Depletion of several factors that control heterochromatin maintenance induces LAD release from the NL (5,6,13,14). However, the casual relationship between heterochromatin homeostasis and its association with the NL has not been understood.

Heterochromatin is thought to be important for genome-NL association because it provides a scaffold of interactions for NE tethers, transmembrane proteins that can physically tether H3K9 methylated regions to the nuclear periphery. A clear example is CEC-4 in C. elegans (15) and LBR (Lamin B receptor) in mammals (16,17), which are considered important tethers of heterochromatin. LBR has been shown to favour conventional genome organisation in metazoan nuclei in synergy with the NL component Lamin A (18).

The transcriptional machinery is an important negative regulator of genome-NL association. Activation of a silent gene associated with the NL promotes physical separation from this component (19–21). Similarly, transcriptional inactivation can promote reattachment (21,22). Conversely, physical detachment of a repressed gene has been shown to allow its activation (23). Furthermore, LAD remodeling during differentiation is coordinated with changes in the transcriptional programme (24–27). With a few exceptions (26), most data suggest a mutually exclusive relationship between gene activation and genome-NL association.

Transcription could shape genome-NL interactions by sequestering active chromatin away from the NL, creating an environment incompatible with NL association. Consistent with this model, in C. elegans the chromatin binding factor MRG1 binds active acetylated regions and sequesters them away from the NL (28). These data suggest a competition model in which different parts of the genome compete for interaction with the NL. This model is also supported by the comparison of genome-NL interactions between haploid and diploid HAP1 cells, which showed a reduction in interaction with the NL for the diploid compared to the haploid haplotype (29). This result supports the idea that there is a limited capacity for chromatin interactions with the NL.

One property of DNA that is emerging as a new level of regulation of chromatin structure and organisation is DNA supercoiling (30,31). This is the ability of DNA or chromatin to change conformation following the application of torsion, stretching or bending of the double helix. These physical forces can arise from the activity of powerful revolving machines such as polymerases, the catalytic activity of chromatin remodelers, or the binding and bending activity of DNA-binding proteins (32,33). Modelling and experimental data suggest that transcriptionally induced supercoiling may contribute to genome folding at multiple levels (31,34,35).

Cells have specialised enzymes to deal with DNA supercoiling, called DNA topoisomerases. These enzymes can catalyse the relaxation of chromatin by introducing transient single (type I topoisomerases) or double (type II topoisomerases) strand breaks on DNA. TOP1, TOP2A and TOP2B are among the most studied topoisomerases and are currently targeted in chemotherapy. While these enzymes have mainly been studied in the context of replication, transcription and cell division (36,37), it has recently been proposed that they may also play a role in genome organisation, particularly in the context of loop extrusion machineries. (38). TOP2B positioning and cutting activity has been described at the base of CTCF-bound chromatin loops in the vicinity of active genes, suggesting that it could regulate the topology between the transcription machinery and the loop-extruding cohesin complex (39–41). In yeast, in interphase, Top2 activity is counteracted by condensin loop extrusion activity and this balance regulates the level of genome entanglement (42). TOP2 activity is also important in regulating mitotic chromosome architecture by regulating sister chromatid resolution and chromatin condensation (43,44). Recently, TOP2 has also been found to act during mitotic exit and its activity is required to reestablish an untangled and compartmentalised genome (45).

Despite the growing evidence that topoisomerases may be involved in 3D genome organisation and folding, the relationship between these enzymes and the anchoring of the genome to the NL has not been investigated. Indeed, the literature on this topic is limited and somewhat sparse. In vitro nuclear reassembly in Xenopus extracts showed that TOP2 inhibitors can block the assembly of a nucleus around protein-free DNA (46,47). Cell-free preparations of Drosophila embryos show that lamins and DNA topoisomerases re-associate on newly assembled nuclei (48). These data suggest a role for TOP2 in the establishment of nuclear architecture. Interestingly, removal of the nuclear envelope reduces segregation defects in topoisomerase mutants (49). TOP2B was found to be important for the transcription of neuronal genes proximal to AT-rich regions that may overlap with lamina-associated domains (50). TOP1 was shown to be able to reduce R-loop formation for long, highly active genes proximal to the NL (51). These data mainly point to a role of the NL as a boundary or constraint.

Apart from this disjointed knowledge on topoisomerases and the NL, a study that addresses whether and how topoisomerases control genome-NL interactions is still missing. In this work, we fill this knowledge gap. We have investigated the role of topoisomerases in genome-NL interactions and linked this to NL tethering. We found that TOP2B is a regulator of genome association with the NL that acts in coordination with the NE tether LBR to maintain proper partitioning of the genome between the NL and the nuclear interior.

## RESULTS

### LADs and iLADs differ in supercoiling state

#### LADs and iLADs have distinct topological states

How DNA supercoiling reflects the anchoring of the genome to the NL has never been investigated. To obtain information on the topological state of LADs in human cells, we used genome-wide maps of DNA supercoiling in RPE1 cells obtained by biotin-psoralen (bTMP) intercalation (52). Psoralen intercalates more frequently in under-twisted DNA (negatively supercoiled) than in relaxed or positively supercoiled DNA (53). In WT RPE1 cells, bTMP-seq revealed a clear differential pattern for iLADs and LADs: iLADs show a higher level of psoralen intercalation. In contrast, LADs show a lower underwound state compared to iLADs (**Figure 1A**). Thus, LADs are less negatively supercoiled than iLADs. Interestingly, on average, LAD borders represented sharp topological transition regions (**Figure 1B**).

**Figure 1.**
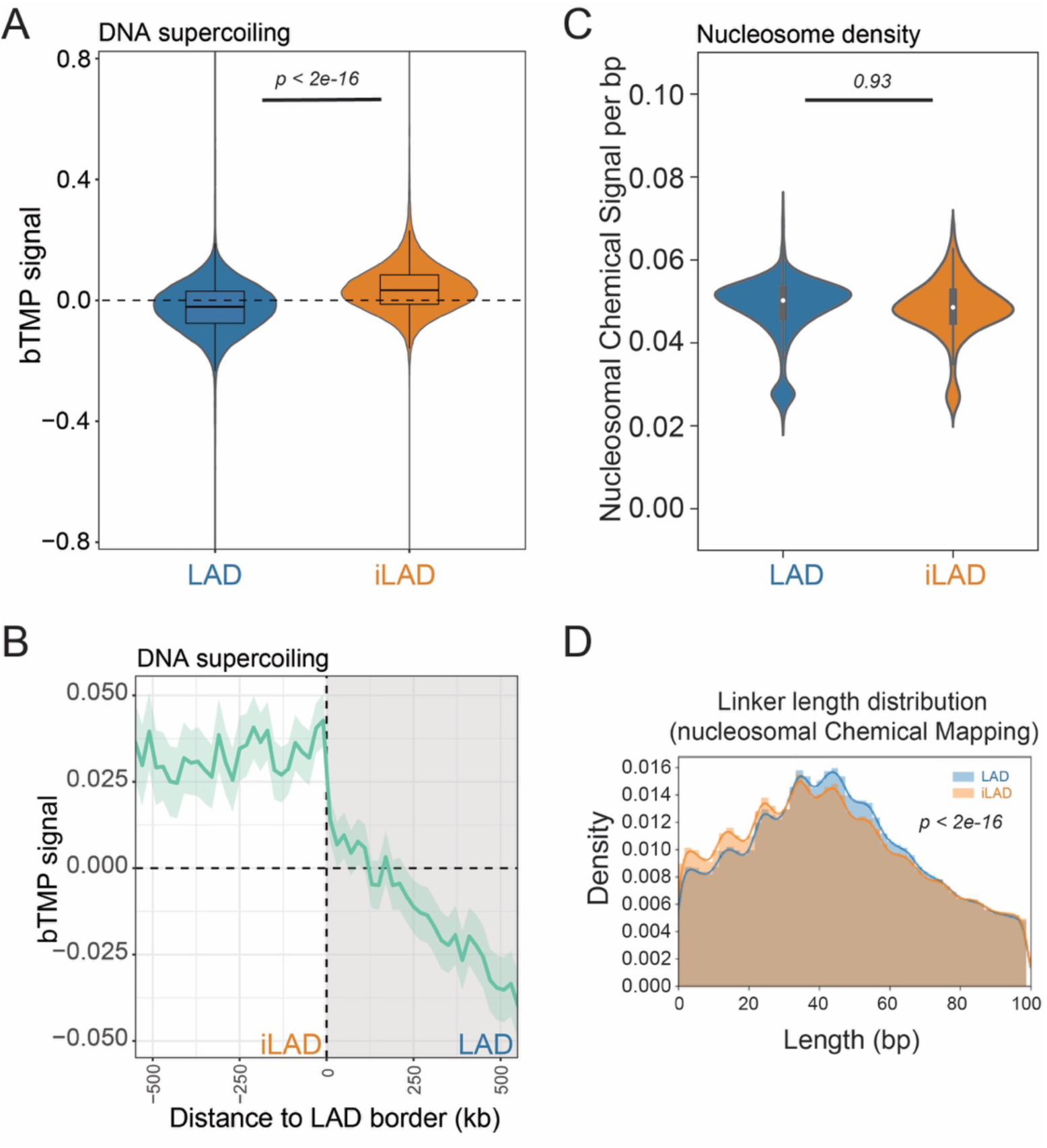
A topological look at LADs. A) bTMP score (log2(IP/input) for LADs and iLADs in WT RPE1 cells. For visualization purposes only, outliers were removed. B) Average bTMP score signal around LAD borders for RPE1 WT cells. The solid line and the shaded area represent the mean signal and 95% confidence interval of the mean, respectively. 500 kb windows inside and outside LADs are shown. For A and B. Results are from two independent biological replicates. (C) Nucleosome signal per base pair for LADs and iLADs in mESC calculated using chemical mapping of nucleosomes. Outlier values from top 0.5 and bottom 0.5 percentiles were excluded from plotting. For A and C, P values are according to Welch’s t-Test. D) DNA linker length distribution from chemical mapping data for LADs and iLADs in mESCs. Data are from (3). Results are from two independent biological replicates. For D, P value is according to Kolmogorov-Smirnov test.

#### Sequence composition does not bias estimates of LAD topology

We considered that a difference in topological state between LADs and iLADs could be due to sequence bias of psoralen intercalation, given that psoralen has a bias for TA dinucleotides (54). However, this sequence preference should lead to increased rather than decreased intercalation in LADs, which are generally AT-rich (55). Furthermore, in our analysis, the psoralen intercalation signal in naked relaxed DNA is subtracted from the bTMP score generated in chromatin, thereby removing any potential DNA sequence bias. Therefore, it is unlikely that the differential supercoiling profile of LAD and iLAD is due to sequence bias.

#### Nucleosome positioning does not bias LAD topology estimates

Nucleosomal occupancy can also reduce psoralen intercalation (56). Thus, the low levels of bTMP intercalation detected in LADs could be explained by a higher nucleosomal density in these regions. To test this, we measured nucleosomal occupancy for LADs and iLADs using MNase data for K562 (from ENCODE, (57)). This analysis showed decreased signal in LADs, suggesting that LADs are depleted of nucleosomes (**Supplementary Figure 1A**). We observed similar results for MNase data generated in mESCs (3) (**Supplementary Figure 1B**). We further explored previously reported maps of nucleosome occupancy in mESCs generated by chemical mapping (3). This revealed similar nucleosome density for LADs and iLADs (**Figure 1C**), suggesting that MNAse-seq may be biased for poor accessibility in LADs. However, linker DNA length distributions calculated from chemical mapping data showed that LADs tend to have longer linker DNA than iLADs (**Figure 1D**). This result suggests that LADs have a slightly lower nucleosomal density than iLADs, which is consistent with data showing that LADs are enriched in histone H1 (58) and that histone H1 binding occurs on longer linker DNA (59). Thus, lower psoralen intercalation in LADs cannot be explained by increased nucleosomal occupancy and likely reflects a topological feature of heterochromatic and NL-bound regions.

Although we cannot completely rule out that other LAD features affecting chromatin accessibility might affect psoralen intercalation in LADs, these data suggest that LADs may have a different topological state than iLADs, with the former being less underwound than the latter.

### TOP2B regulates LAD partitioning *in vivo*

#### TOP2B controls the partitioning of the genome between LADs and iLADs

To understand the impact of DNA topology on genome-NL association *in vivo*, we depleted either TOP1, TOP2A or TOP2B by RNA interference and mapped genome-NL interactions using pA-DamID, a recent adaptation of DamID that provides improved temporal resolution (60). (**Supplementary Figure 1C**). Interestingly, all three DNA topoisomerases could modulate genome-NL interactions to some extent and at specific genomic locations, with both gain and loss of DNA-NL interactions following topoisomerase loss (**Figure 2A**). However, we observed a more robust effect of TOP2B depletion. Unlike TOP1 and TOP2A, TOP2B depletion led to a substantial redistribution of genome-NL contacts. This was characterised by a switch from the bimodal distribution typical of control cells to a monomodal distribution in TOP2B-depleted cells (**Figure 2B**), implying an overall loss of partitioning of the genome between the NL and the nuclear interior. Gains and losses of genome-NL interactions involved large parts of the genome, and changes could be detected in both LAD and iLAD regions (**Figure 2A**).

**Figure 2.**
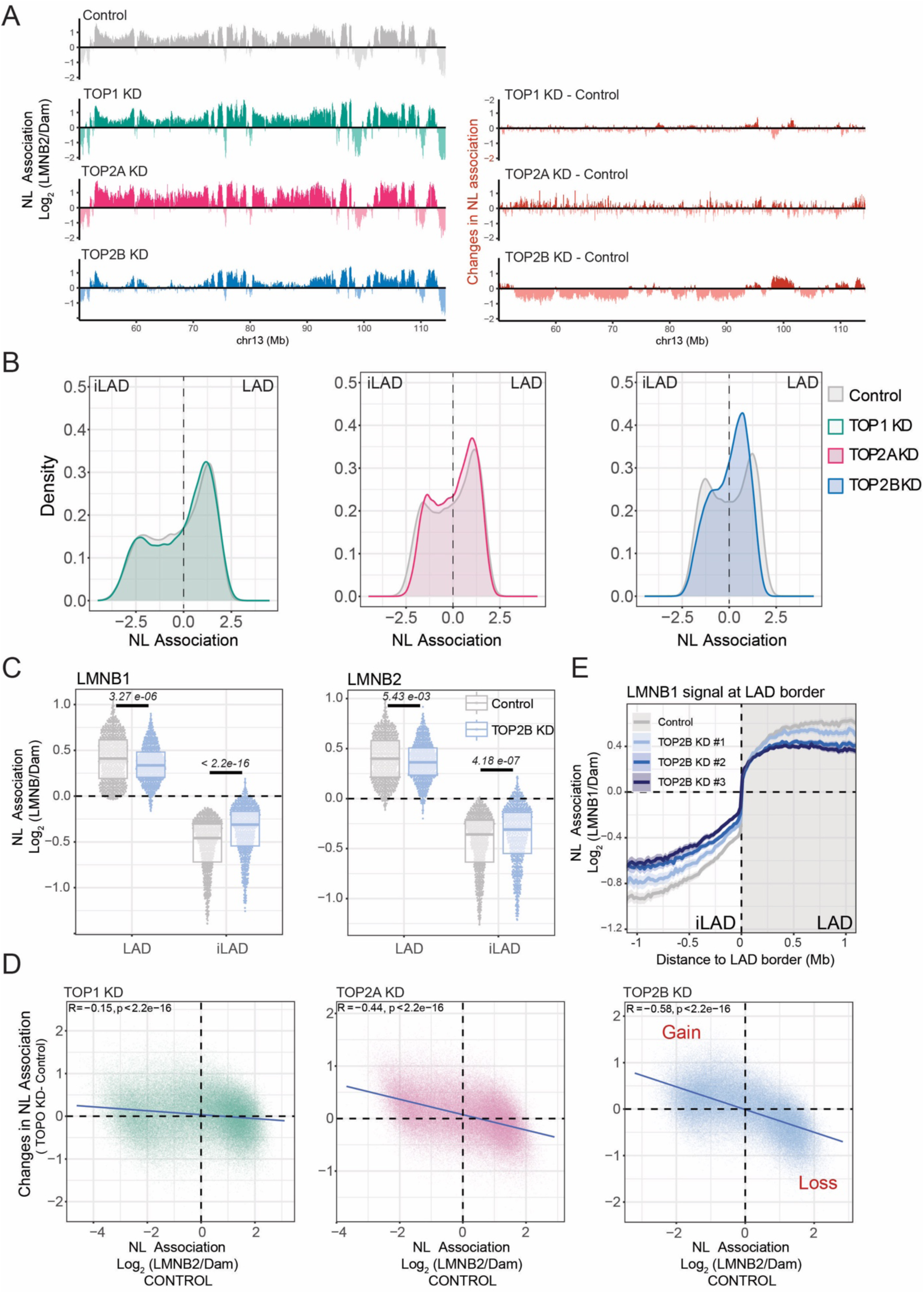
TOP2B controls LAD partitioning. A) *Left* Representative genomic track of LMNB2 pA-DamID for control sample (grey), TOP1 (green), TOP2A (purple), and TOP2B (blue) knockdowns. 20-kb bins were used. The antibody signal is normalized over a Damonly control. *Right:* Differential tracks (knockdown - control, orange) highlighting the gain and loss of genome-NL interactions following the depletions of the three topoisomerases. B) LMNB2 signal distribution for control, TOP1, TOP2A, and TOP2B depletion. In control cells, a bimodal distribution usually represents iLAD/LAD partitioning. Results are from three independent biological replicates. C) LAD and iLAD score following TOP2B depletion using LMNB1 and LMNB2 mapping data. Results are from two biological replicates and 4 technical replicates by using two control siRNAs and two TOP2B-specific siRNAs. D) Correlation scatter plots of 20 kb genomic bins for LMNB2 score in control cells (x-axis) and differential LMNB2 score (topoisomerase depletion - control, y-axis) for TOP1, TOP2A, and TOP2B depletions. The blue line represents a linear model, Pearson correlation and *p*-value are shown in the plots. E) Average LMNB1 pA-DamID score around LAD borders for control and TOP2B depleted samples. The effect of three different siRNAs for TOP2B depletion are shown separately. Genome-NL contact levels in control samples are the average of two biological replicates and 4 technical replicates generated using two control siRNAs. The solid line and the shaded area represent the mean signal and 95% confidence interval of the mean, respectively.

Following TOP2B depletion, LADs showed a significant reduction in NL association and iLADs often gained NL interactions. This trend was observed by pA-DamID of both LMNB1 and LMNB2 (**Figure 2C**, **Supplementary Figure 1D**). Such a de-partitioning effect could be adequately described as the slope in a correlation plot of the NL association score in control cells versus changes in NL interactions following topoisomerase depletion (**Figure 2D**). This analysis revealed substantially stronger effects for TOP2B loss than for the other two topoisomerases. Although TOP2A knockdown had some effects on genome-NL interactions, we decided not to investigate this enzyme further due to the apparent strong effects on cell cycle progression, consistent with the known role of TOP2A in decatenation and progression through mitosis {Uuskula-Reimand, 2022, 36322662). In contrast, TOP2B depletion did not alter cell cycle progression (**Supplementary Figure 1E**).

#### Changes induced by TOP2B depletion are robust and reproducible

To verify these results, we performed several additional controls. First, we induced TOP2B depletion with two additional siRNAs. All depletion experiments showed similar effects and a high degree of genome-wide correlation, ruling out siRNA-specific effects (**Figure 2E, Supplementary Figures 2A and 2B**). Second, we considered that the pA-DamID score is a normalised signal calculated as the log_2_(ratio) of two different signals, one from an antibody (e.g. LMNB2) and the other from the use of a freely diffusible Dam. This second signal takes into account potential changes in DNA accessibility or sequence distortion. To rule out the possibility that TOP2B loss causes changes in genome accessibility, we examined the changes in the ‘Dam only’ and ‘LMNB2 only’ datasets after TOP2B knockdown. We found that most of the normalised signal changes were due to changes in LMNB2 levels (**Supplementary Figures 3A and 3B**). Third, to rule out a possible normalisation problem, we performed calibrated pA-DamID in RPE1 cells (control and TOP2B depleted) spiked in with mouse embryonic fibroblasts (**Supplementary Figures 3C**, see **Methods**). Data calibrated to mouse reads showed genome-wide profiles almost identical to uncalibrated data for changes in chromatin/NL contacts after TOP2B knockdown (**Supplementary Figures 3D and 3E**). Thus, the changes in genome-NL interaction after TOP2B depletion described here are not an artefact of the normalisation process.

#### TOP2B control over genome-NL interaction is conserved in other cell lines

Finally, we extended our analysis to additional cell lines by measuring genome-NL interactions in HCT116 and K562 cell lines that are hypomorphic for TOP2B. Both cell lines showed weaker LAD/iLAD partitioning upon TOP2 depletion, although to a lesser extent than in RPE1 cells (**Supplementary Figures 4A-D**). Taken together, these data support a role for TOP2B in the control of genome-NL interactions.

### Changes in genome-NL interactions after TOP2B depletion are not explained by changes in transcription

#### Transcription is relatively stable during TOP2 KD

We and others have previously shown that transcriptional activation promotes the physical separation of chromatin from the NL {Tumbar, 2001, 11175745; Therizol, 2014, 25477464; Brueckner, 2020, 32080885}. Similarly, transcriptional inhibition favours genome-NL reassociation (21,22). Since TOP2B is considered to be a transcription-associated topoisomerase (61), we asked whether changes in gene expression following TOP2B depletion could explain changes in genome-NL interactions. RNA-seq analysis revealed modest changes in transcription upon TOP2B depletion, with 917 genes significantly upregulated and 645 genes downregulated (**Supplementary Figure 5A**). However, the overall disregulation was relatively marginal, as only 219 upregulated and 118 downregulated genes showed more than 2-fold change, suggesting that TOP2B depletion did not induce dramatic changes in gene expression at the time point analysed. Differentially expressed genes showed significant but very modest changes in the pA-DamID score (**Supplementary Figure 5B**). Similarly, genes with a differential DamID score showed significant but marginal changes in expression levels (**Supplementary Figure 5C**). These changes could explain only a fraction of the overall alterations in NL association observed after TOP2B depletion. Thus, it is unlikely that transcriptional changes induced by TOP2B knockdown are the main cause of the pronounced de-partitioning phenotype.

### Dissipation of dynamic supercoiling is not sufficient to promote LAD reshaping

#### In vivo chromatin relaxation by break inducers does not alter genome-NL interactions

Two types of DNA supercoiling can be identified in chromatin: unconstrained supercoiling generated by active processes such as transcription or replication and constrained supercoiling typical of DNA-protein interactions (32,33). If LAD partitioning is due to simple dynamic supercoiling, then its dissipation by single-strand break inducers should be sufficient to promote some changes in genome-NL contact. To test this hypothesis, we performed *in vivo* chromatin relaxation experiments: we mapped genome-NL interactions after three hours of treatment with two doses of bleomycin. This drug can induce single and double-strand breaks in a dose-dependent manner, promoting the dissipation of dynamic, unrestrained supercoils (52) by leaving restrained supercoils intact. At the tested concentrations and time, bleomycin treatment did not affect genome partitioning at the NL. (**Supplementary Figure 6A**). Although we cannot rule out that genome-NL interactions could change at different time points during bleomycin treatment, these data suggest that simple dissipation of dynamic supercoils induced by DNA break inducers is not sufficient to promote changes in genome-NL interactions.

#### Ex vivo chromatin relaxation by purified topoisomerases does not alter genome-NL interactions

DNA topoisomerase activity can remove supercoils from chromatin. We performed *ex vivo* chromatin relaxation by treating permeabilized nuclei with purified human type II topoisomerases, either TOP2A or TOP2B. After 30 minutes of incubation in the TOP2 relaxation buffer in the presence or absence of hTOP2A or hTOP2B, permeabilized nuclei were processed for classical pA-DamID protocol for LMNB2. We detected minor differential genome-NL interactions after incubation with topoisomerase II as compared to incubation with the TOP2 buffer alone (**Supplementary Figure 6B**). Although we cannot rule out that exogenous TOP2 proteins were unable to sufficiently access the genomic DNA under these conditions, these results suggest that chromatin relaxation and/or decatenation alone are not sufficient to alter genome-NL interactions *ex vivo*.

### TOP2B might protect iLADs from genome-NL association

#### TOP2B positioning and activity are mainly enriched in iLADs

To better understand the relationship between TOP2B and genome-NL association, we focused on TOP2B positioning in the genome. We generated pA-DamID data for TOP2B and validated this in our TOP2B-depleted cells (**Figure 3A**). The mapping data showed that TOP2B is relatively depleted in LADs and mainly enriched in inter-LAD locations, although we could also detect a low TOP2B-specific signal in LADs (**Figure 3A and 3B**, **Supplementary Figure 7A**).

**Figure 3.**
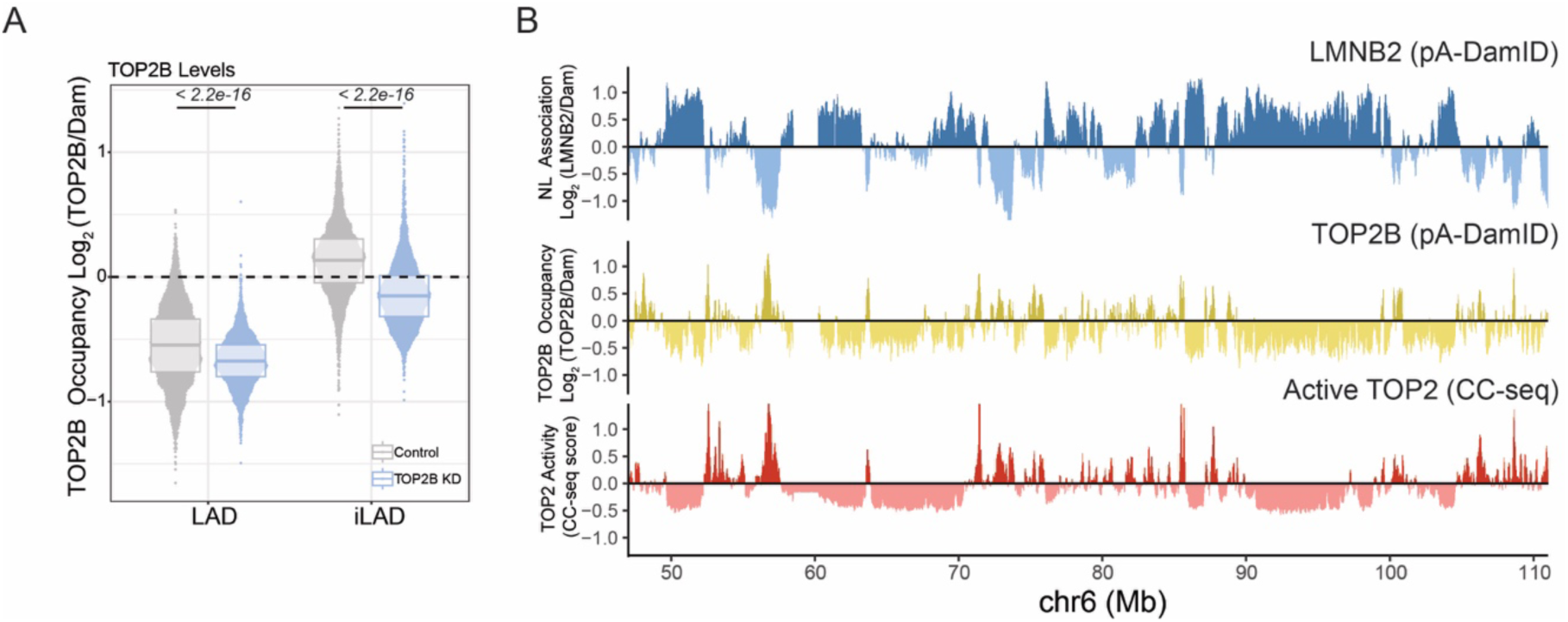
Top2 positioning and activity relative to LADs. A) TOP2B pA-DamID score in LAD and iLAD in control and TOP2B depleted cells. Results are from three independent biological replicates. P values are according to Wilcoxon’s test. B) Representative genomic tracks for pA-DamID signal for LMNB2 (blue) and TOP2B (yellow) and CC-seq (red). CC-seq data are from (1) and are from at least three biological replicates.

Next, we focused on the localisation of catalytically active TOP2B. We used mapping data of catalytically active TOP2 in RPE1 cells generated by CC-seq (1). This technique detects cleavage complex formation when TOP2 is poisoned on DNA by VP16 treatment, and is a measure of catalytically engaged TOP2. Although CC-seq cannot distinguish between TOP2A and TOP2B, in RPE1 cells most of the CC-seq signal is derived from TOP2B activity (1). Therefore, this technique can be used as a proxy for TOP2B activity. We found a high positive correlation between our TOP2B pA-DamID data and the CC-seq dataset (**Figure 3A**, **Supplementary Figure 7B**). Similar to our pA-DamID data, CC-seq showed that LADs are relatively depleted of high TOP2 activity compared to iLADs (**Supplementary Figure 7C**). Considering that iLADs tends to show a gain of interactions with the NL upon topoisomerase depletion (**Figure 2C-D**) these data might suggest that TOP2B might protect iLADs from association to NL.

### A TOP2B/LBR axis controls genome partitioning at the NL

#### Mapping of LBR-associated chromatin

Our data suggest that TOP2B may protect iLADs from interacting with the NL. However, this model does not fully explain why genome-NL interactions in LADs become weaker following TOP2B depletion. Indeed, a competition model would require LADs to be released from the NL. We wondered whether TOP2B could mediate genome-NL interactions in coordination with an NE tether. We focused on the lamin B receptor, LBR. This protein is an organiser of heterochromatin at the nuclear periphery (62,63). Interestingly, in vitro data suggest that the chromatin-interacting nucleoplasmic domain of LBR binds DNA with an affinity dependent on DNA supercoiling (64). These considerations led us to investigate a possible link between TOP2B and LBR. We first mapped contacts between the genome and LBR utilizing pA-DamID and an LBR-specific antibody. We confirmed the high specificity of the resulting maps, by using an LBR knockout cell line as control (**Supplementary Figure 8A**). In wild-type cells, genome-LBR interactions correlated strongly with genome-LMNB2 interactions (**Supplementary Figure 8B**) and anti-correlated with TOP2 activity (**Supplementary Figure 8C**), in agreement with previous data (**Supplementary Figure 7C**). In general, LBR mapping showed a partitioning effect similar to mapping of LMNB1 and LMNB2 interactions following TOP2B depletion (**Supplementary Figure 8D**), and the changes induced by TOP2B loss on genome-LBR interactions correlated strongly with changes in genome-LBR contacts (**Supplementary Figure 8E**).

#### A subset of LADs preferentially lose contact with LBR following TOP2B depletion

A direct comparison of LMNB1, LMNB2, and LBR at the LAD level revealed a more pronounced and significant weakening of interactions of LADs with LBR (**Figure 4A, Supplementary Figure 8F**). This result was particularly striking considering that the LBR signal in RPE1 cells had a significantly lower dynamic range compared to signals generated with LMNB1 and LMNB2 mapping and should therefore be less sensitive in capturing potential differences (**Supplementary Figure 8G**). A visual comparison of differential z-scores (TOP2BKD-Control) for the three markers revealed a detachment detectable in LAD regions, significantly stronger for LBR-chromatin contacts (**Figure 4B**). Consistent with these observations, Limma-Voom differential analysis (60,65) identified 108 LADs (out of 765 LADs, 14%) that significantly lose interactions after TOP2B depletion, specifically with LBR, but not with LMNB1 and LMNB2 (**Figure 4C**). We conclude that in response to TOP2B loss, a subset of LADs preferentially lose contact with the nuclear envelope tether LBR.

**Figure 4.**
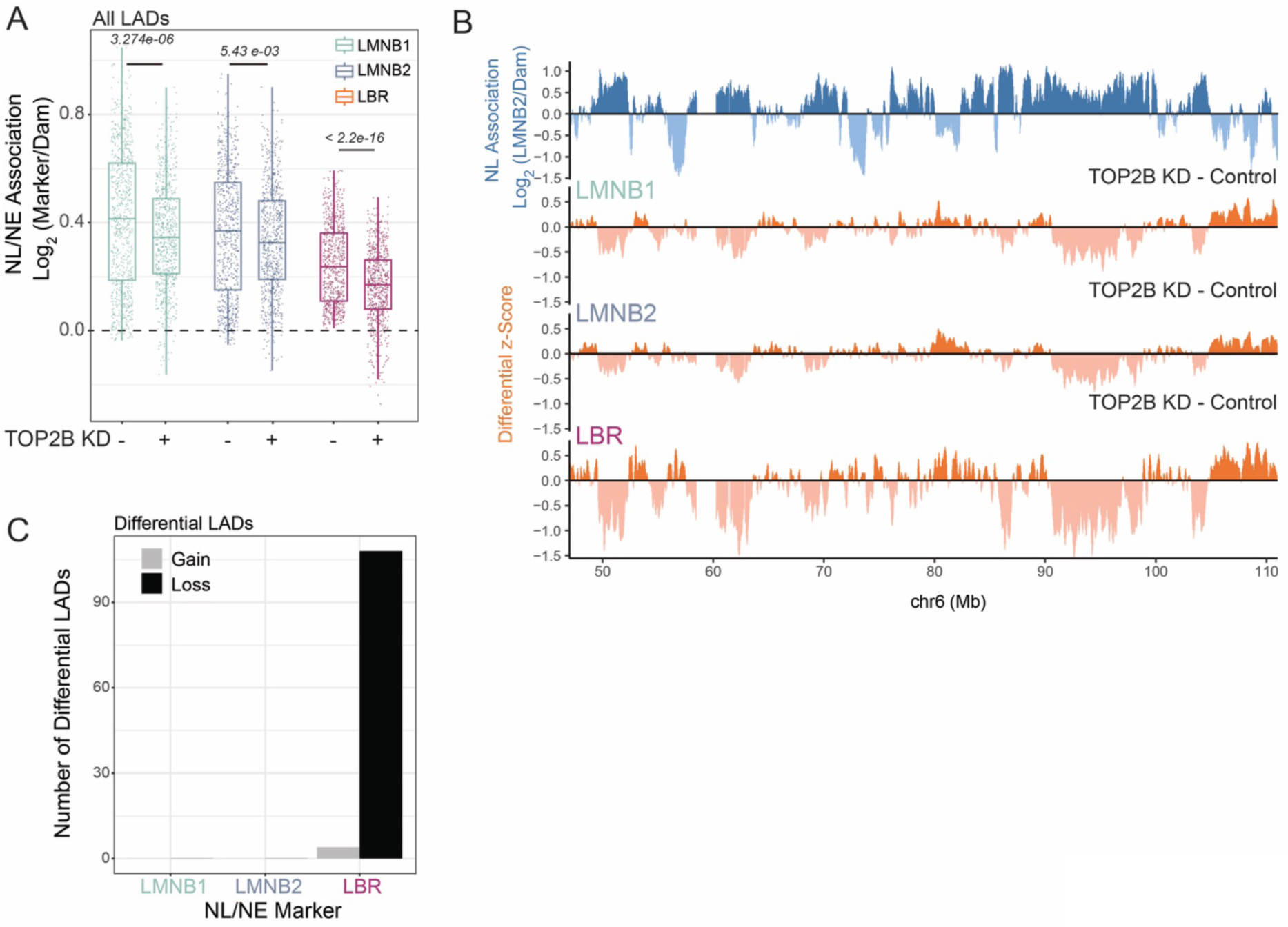
A subset of LADs preferentially loses contacts from LBR following TOP2B loss. A) LAD pA-DamID scores for control and TOP2B depleted cells for three different marks: LMNB1, LMNB2 and LBR. P values are according to Wilcoxon’s test. B) Representative genomic tracks of LMNB2 signal (pA-DamID, blue) for control RPE1 cells and relative differential tracks (TOP2B knockdown – control, orange) using z-scored signal from three different antibodies: LMNB1, LMNB2 and LBR. 20-kb bins were used. One unit of z-score corresponds to 0.297, 0.204, 0.219 in differential pA-DamID score for LMNB1, LMNB2 and LBR, rispectively. C) Results of Voom-Limma analysis used to call LADs that significantly gain or lose interaction with LMNB1, LMNB2 and LBR following TOP2B depletion. Results are from two biological replicates and 4 technical replicates generated using two negative control siRNAs and two TOP2B-specific siRNAs.

#### LBR knockout causes substantial changes in genome-NL interactions

To investigate a possible link between TOP2B and LBR, we generated LMNB2 pA-DamID data in RPE1 cells in which we knocked out LBR by CRISPR genome editing (**Supplementary Figure 8H**, (23)). In RPE1 the loss of LBR caused a strong de-partitioning of genome-NL interactions (**Supplementary Figure 8I**) and profound changes in LAD strength, with strong LADs showing a marked reduction of LMNB2 signal (**Supplementary Figure 8L**). Differential LAD calling analysis identified 220 LADs significantly losing interactions with LMNB2. Three hundred LADs increased their NL association upon LBR loss, a phenomenon we did not observe upon TOP2B depletion. This result probably reflects compensatory movements induced by the prolonged absence of LBR.

#### TOP2B knockdown mirrors the effects of LBR knockout

A direct comparison with regions that lose contact with LBR upon TOP2B depletion revealed that approximately 87% of LADs that significantly detach from LBR following TOP2B knockdown also undergo NL detachment upon LBR knockout (**Figure 5A**). Similarly, almost half of the LADs that significantly lose interaction with the NL in LBR knockout cells also detach upon TOP2B depletion. These data suggest that TOP2B and LBR may modulate genome-NL contacts in a similar manner. To test this directly, we compared the two differential scores (perturbation - control) for TOP2B knockdown and LBR knockout. We found a striking correlation between changes in genome-NL contacts after TOP2B depletion and after LBR knockout (**Figure 5B and 5C**). The magnitude of the effect was greater for LBR knockout than for TOP2B depletion. These data show that TOP2B and LBR control the interactions between the genome and the NL in very similar ways. It is also important to note that loss of TOP2B did not induce changes in either transcript (Log_2_(fold change) = −0.21, p adj =0.54) or protein levels for LBR (**Supplementary Figure 8H**).

**Figure 5.**
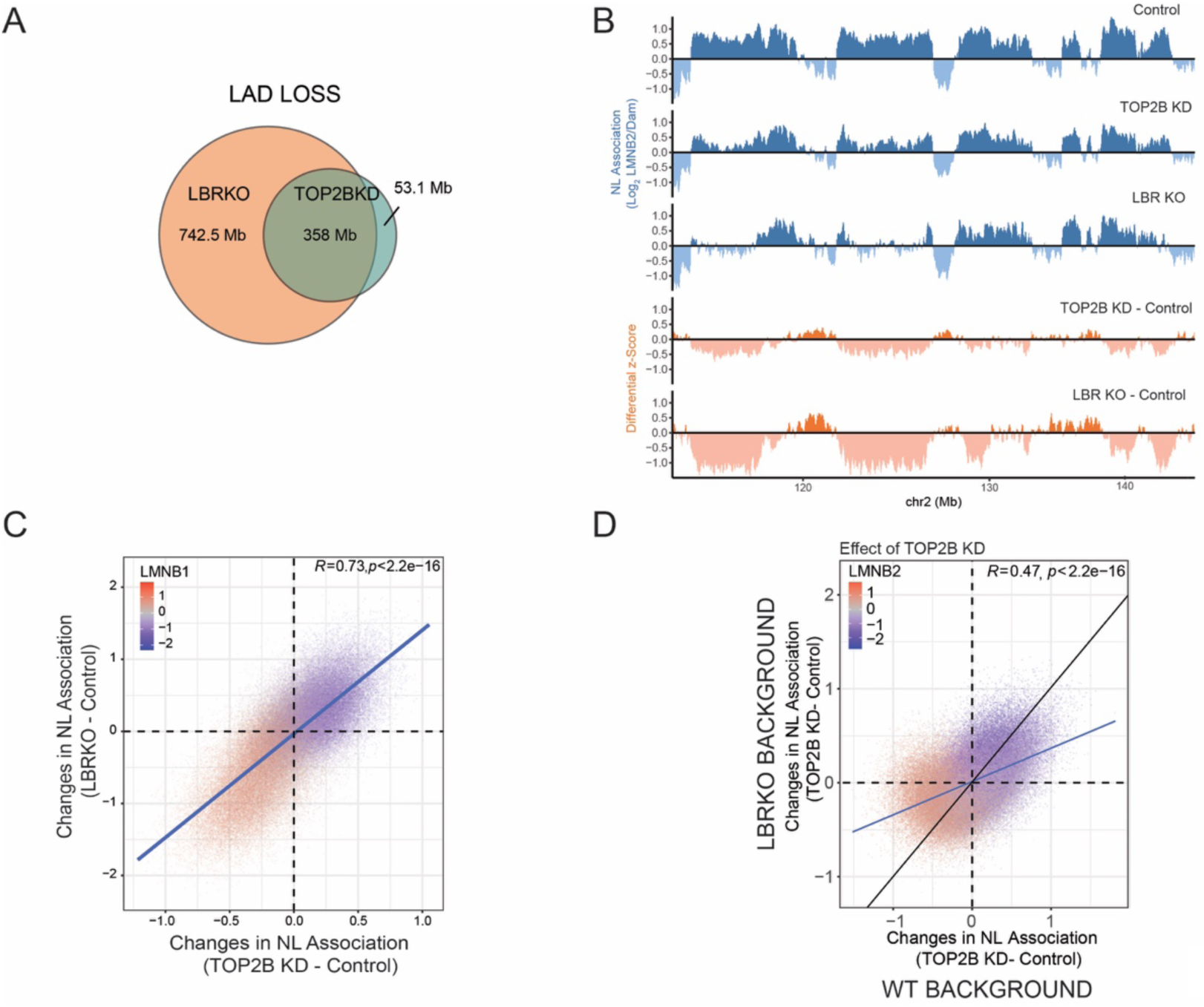
A TOP2B/LBR axis controls genome partitioning at the NL. A) Genomic overlap (in Mb) for LADs released by LBR during TOP2B depletion and LADs that significantly lose genome-NL interactions following LBR Knockout. B) *Blue tracks:* representative genomic tracks of LMNB2 pA-DamID for the control sample, TOP2B knockdown, and LBR knockout. 20-kb bins were used. *Orange tracks:* differential z-scored LMNB2 tracks (depletion-control) for TOP2B and LBR depletions. C) Correlation scatter plot for differential LMNB2 pA-DamID scores for TOP2B depletion (TOP2B knockdown - control, x-axis) and LBR knockout (y-axis LBR knockout - control, y-axis). The results are an average of 4 biological replicates for LBR knockout. For TOP2B depletion results are from two biological replicates and 4 technical replicates by using two control siRNAs and two TOP2B specific siRNAs. D) Correlation between changes in genome-NL interactions induced by depletion of TOP2B in WT and LBR knockout background. Results are from 4 biological replicates.

#### Co-depletion points to a sub-additive role for TOP2B and LBR

To better dissect the role of TOP2B and LBR in genome partitioning to the NL, we depleted TOP2B in the LBR knockout background (**Supplementary Figure 8H**). In the absence of LBR, depletion of TOP2B caused further de-partitioning, mostly of the same genomic regions as it controlled in the presence of LBR (**Figure 5D**). However, we observed stronger changes in the NL interaction in the presence rather than in the absence of LBR, as shown by a flattening of the correlation line between the differential scores in the two different genetic backgrounds (**Figure 5D**). Thus, TOP2B and LBR are neither epistatic nor synergistic in their effects on NL interactions, but show sub-additive phenotypes.

### Co-depletion of TOP2B and LBR recapitulates LAD reshaping typical of oncogene-induced senescence

#### TOP2B and LBR depletion mimics changes typical of OIS

The co-depletion of TOP2B and LBR showed the most substantial de-partitioning effect with genomic regions switching from LAD to iLAD state and vice versa (**Figure 6A and Supplementary Figure 9B**). This relatively strong phenotype reminded us of strong LAD reshaping events typical of oncogene-induced senescence (OIS). During this type of senescence, constitutive LADs (cLADs), defined as LADs conserved among different cell lines, detach from the NL (4,66). We wondered if LADs detaching after TOP2B depletion were predominantly cLADs. Using a LAD atlas that identified constitutive and facultative LADs (29) (fLADs, LADs that are not maintained across different cell lines), we found an overlap between lost LADs induced by TOP2B knock-down and constitutive LADs, significantly higher than the overlap with fLADs and constitutive inter-LADs (ciLADs) (**Figure 6B**). Thus, similarly to what happens during OIS, constitutive LADs become weaker following TOP2B depletion.

**Figure 6.**
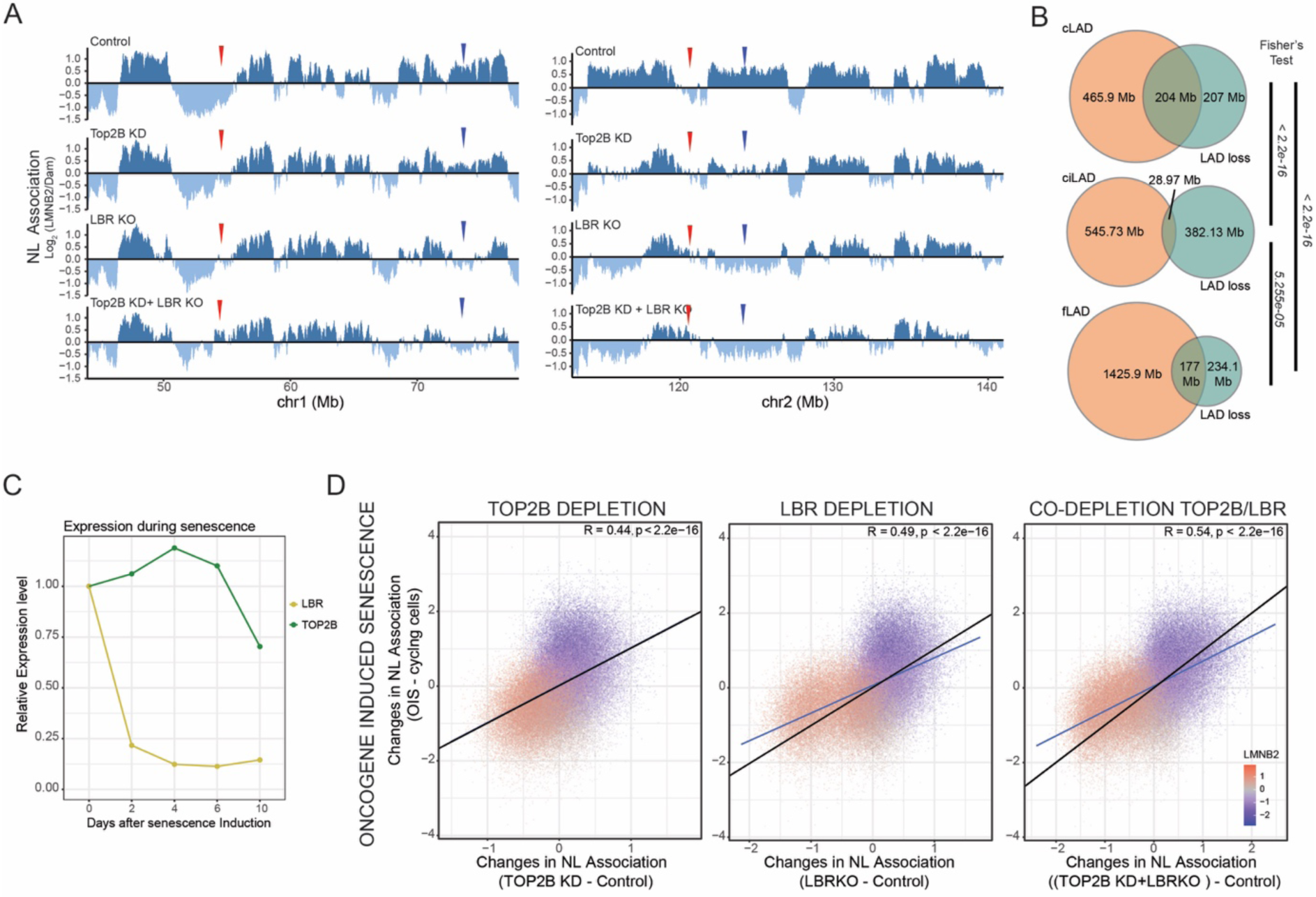
TOP2B and LBR co-depletion mimics LAD reshaping typical of oncogene-induced senescence. A) *Partial LAD/iLAD switch:* Two representative genomic tracks of LMNB2 pA-DamID for the control sample, TOP2B knockdown, LBR knockout, and co-depletion of TOP2B and LBR. 20-kb bins were used. The antibody signal is normalized over a Dam-only control. *Red arrows*: regions that progressively switch from iLAD to LAD state during the different depletions. *Black arrows*: regions that progressively switch from LAD to iLAD state during the different depletions. B) Genomic overlap in Mb for LADs released by LBR during TOP2B depletion and constitutive LADs, facultative LADs and constitutive iLADs. C) RNA-seq data showing transcript level for LBR and TOP2B during OIS. Results are from (2) and are the average of three independent biological replicates. D) Correlation scatter plot of 20-kb genomic bins for differential LMNB2 scores (x-axis) for TOP2B knockdown (left plot), LBR knockout (middle plot), and codepletion of TOP2B and LBR (left plot) and changes in lamina association after OIS (y-axis). The blue line represents a linear model; Pearson correlation and *p*-value are shown in the plots. Data in RPE1 cells are. The average of 4 biological replicates. OIS data are from (4) and are the average of three independent biological replicates. Since OIS data were generated in Tig3 fibroblasts and not RPE1 cells, discordant bins in control data between the two cell lines were filtered out to allow direct comparisons of conserved genome-NL interactions. This led to the removal of near 30% of the total number of 20-Kb bins.

This prompted us to examine the expression of TOP2B and LBR during OIS using published RNA-seq data (2). Interestingly, while LBR expression is downregulated during OIS, TOP2B expression remains relatively high, suggesting that these two proteins are differentially regulated during OIS (**Figure 6C**). To further explore a possible link between TOP2B, LBR and senescence, we directly compared our depletion experiments (TOP2B knockdown, LBR knockout and co-depletion) with DamID data generated in OIS cells (4). We found positive correlations between changes in genome-NL interactions during all perturbations and those typical of OIS. The correlation was particularly robust when we compared OIS with co-depletion of TOP2B and LBR (**Figure 6D**). Cell cycle analysis revealed no alteration in cell cycle progression for either single or double depletions (**Supplementary Figures 1D and 9B**). Thus, despite mimicking the LAD remodelling typical of OIS, single and co-depletion of TOP2B and LBR did not induce senescence.

### Heterochromatin is stable in detached LADs during TOP2B depletion

#### LAD heterochromatin marked by H3K9me3 is repositioned after TOP2B loss

LADs are heterochromatic regions frequently marked by H3K9 and H3K27 methylation (5,8,67), and LBR is well known as a tether that anchors heterochromatin at the NE (16,17,62). In addition, data suggest that TOP2 can regulate heterochromatin in both yeast and mammals (68,69). We investigated whether heterochromatin in LADs is stable following loss of TOP2B and partial release from LBR by using pA-DamID to map H3K9me3, H3K9me2 and H3K27me3 genome-wide. In control RPE1 cells, H3K9me3 was enriched in the central part of the LADs. H3K9me2 and H3K27me3 marked the distal part of LAD regions as previously described (70,71) (**Supplementary Figure 10A**), and were relatively less abundant in the part of LADs where H3K9me3 was present. Notably, changes in genome-NL interactions after TOP2B depletion were negatively correlated with H3K9me3, but not with H3K9me2 and H3K27me3 (**Figure 7A and 7B**). This result suggests that strong LADs that detach from the NL after TOP2BKD are enriched in H3K9me3 and point to a mechanism where this type of heterochromatin is released from LBR after TOP2B depletion. Thus, in RPE1 cells, TOP2B controls the distribution of H3K9me3-marked heterochromatin at the NE.

**Figure 7.**
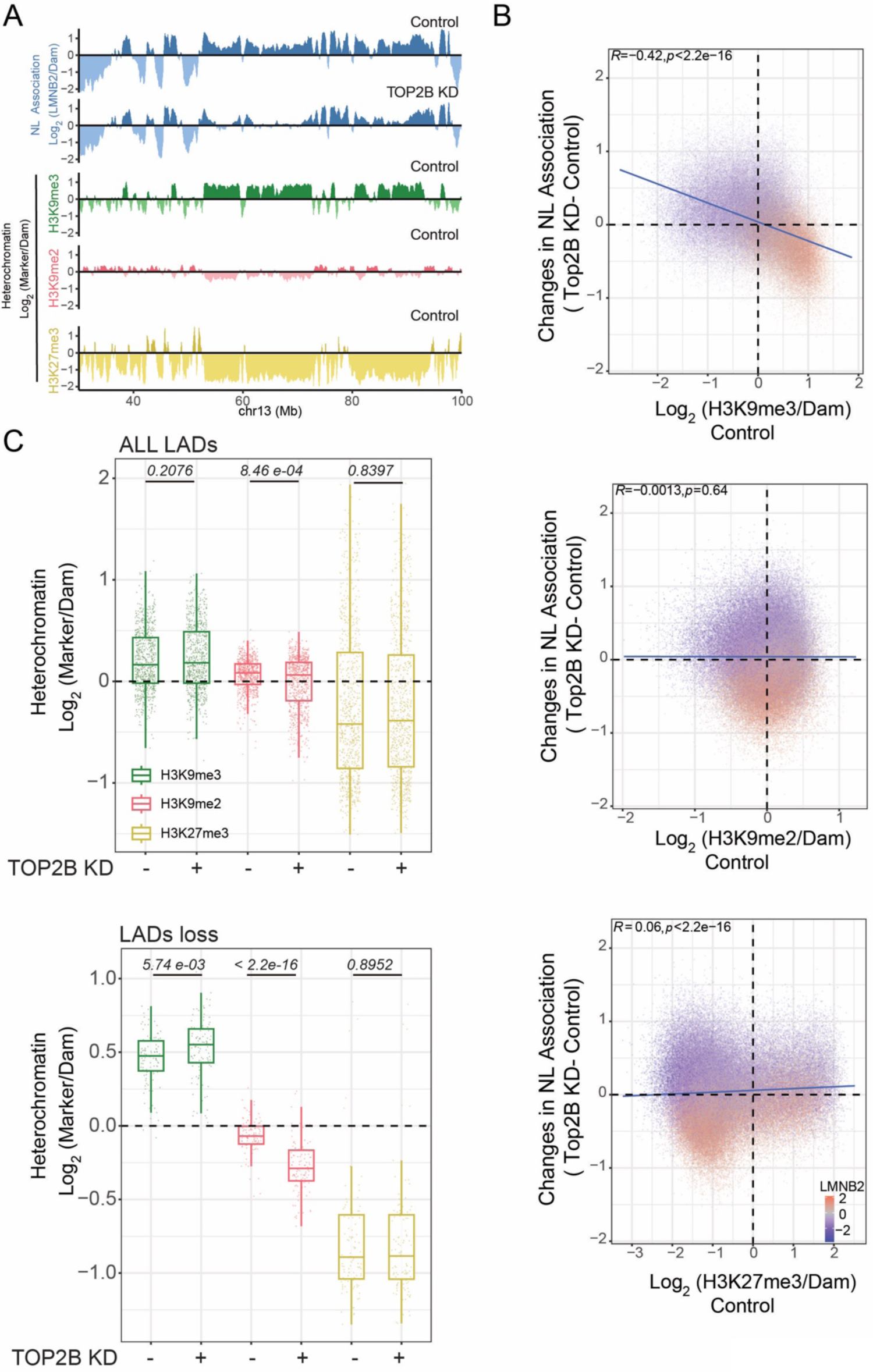
Stability of heterochromatin in LADs released by LBR after TOP2 loss. A) Representative genomic track for pADamID signal for LMNB2 (blue, control and TOP2BKD cells), H3K9me3 (green, control cells), H3K9me2 (red, control cells), and H3K27me3 (yellow, control cells). 20-kb bins were used. The antibody signal is normalized over a Dam-only control. B) *Constitutive heterochromatin (H3K9me3) detaches from the NL during TOP2BKD*. Correlation scatter plot of 20-kb genomic bins for H3K9me3, H3K9me2, and H3K27me3 scores in control cells (x-axis) and differential LMNB2 score (TOP2BKD - control, y-axis). The blue line represents a linear model; Pearson correlation and *p*-value are shown in the plot. C) H3K9me3, H3K9me2, and H3K27me3 scores in control and TOP2B depleted cells in all LADs (top panel) and LADs released by LBR after TOP2B knockdown (bottom panel). Results are from two independent biological replicates. P values are according to Wilcoxon’s test.

#### Stability of H3K9me3 following TOP2B loss

We then measured the levels of H3K9me3, H3K9me2 and H3K27me3 after TOP2B depletion. We calculated a heterochromatin score of the three marks for LADs released from LBR following TOP2B knockdown and for the ensemble of all LADs as a control. The data show that the levels of H3K9me3 increased slightly in regions released from LBR following TOP2B loss.Thus, H3K9me3 is not lost following TOPB depletion, but rather slightly enhanced following release from the LBR (**Figure 7C, Supplementary Figure 10B**). These dynamics may reflect a compensatory mechanism triggered by release from the NL.

#### H3K9me2 is sensitive to TOP2B loss

We noticed that H3K9me2 showed a peculiar behaviour. Although this mark is relatively low in LADs that lose contact with LBR, its modulation after TOP2B KD follows the changes in DNA-NL association. LADs that preferentially lost interactions with LBR lost even more H3K9me2 (**Figure 7C, Supplementary Figure 10B**). We propose that H3K9me2 is sensitive to the NL environment. However, given the low levels of this mark in these regions, it is unlikely that these changes are the drivers of the genome-NL interactions that are reshaped during TOP2B depletion. Thus, these data show that following TOP2B depletion, H3K9me3-marked heterochromatin released from LBR is not lost.

## DISCUSSION

This study reveals a role for TOP2B in regulating chromatin contacts with the NL. Upon TOP2B depletion, constitutive LADs marked by H3K9me3 preferentially detach from the NL, whereas iLADs show increased NL interactions. Importantly, H3K9me3 itself remains largely unaffected. These data suggest a mechanism by which heterochromatin is not disrupted during TOP2B knockdown and is instead repositioned away from the NL. These changes in NL interactions are remarkably similar to those observed during oncogene-induced senescence (4) (**Figure 6**).

Several lines of data in C. elegans and humans have suggested a competition model for genome partitioning at the NL. This model proposes that in a single nucleus, the NL has a limited capacity to interact with the genome (5,21,28,29,72). We found that the catalytic activity of TOP2B is mainly concentrated in iLAD regions, and that some of these regions gain interactions with the NL following topoisomerase depletion. These data suggest that topological control of iLADs by TOP2B may protect this part of the genome from association with the NL. If TOP2B is limiting, iLADs might interact more frequently with the NL and compete with LADs for NL association. This phenomenon could be enhanced by the concomitant release of LADs when LBR is simultaneously depleted.

We propose that the activity of TOP2B and LBR may help to compartmentalise the genome between the nuclear periphery and the nuclear interior. Mapping data show that these two proteins occupy two distinct genomic compartments in interphase: LBR interacts with LADs and TOP2B binds to iLADs. Our co-depletion experiments show that TOP2B depletion can still trigger changes in genome-NL interactions even without LBR, suggesting that these two proteins are not epistatic. We propose that these two proteins form an axis that allows correct genome segregation between the NL and the interior by acting on two opposite parts of the genome (**Figure 8A**). The striking similarity in the way LBR and TOP2B shape genome-NL interactions could suggest that the activity of these two proteins is linked and coordinated. This hypothesis is supported by the more pronounced dissociation of LAD chromatin from LBR than from lamins upon loss of TOP2B. The underlying mechanism remains to be elucidated.

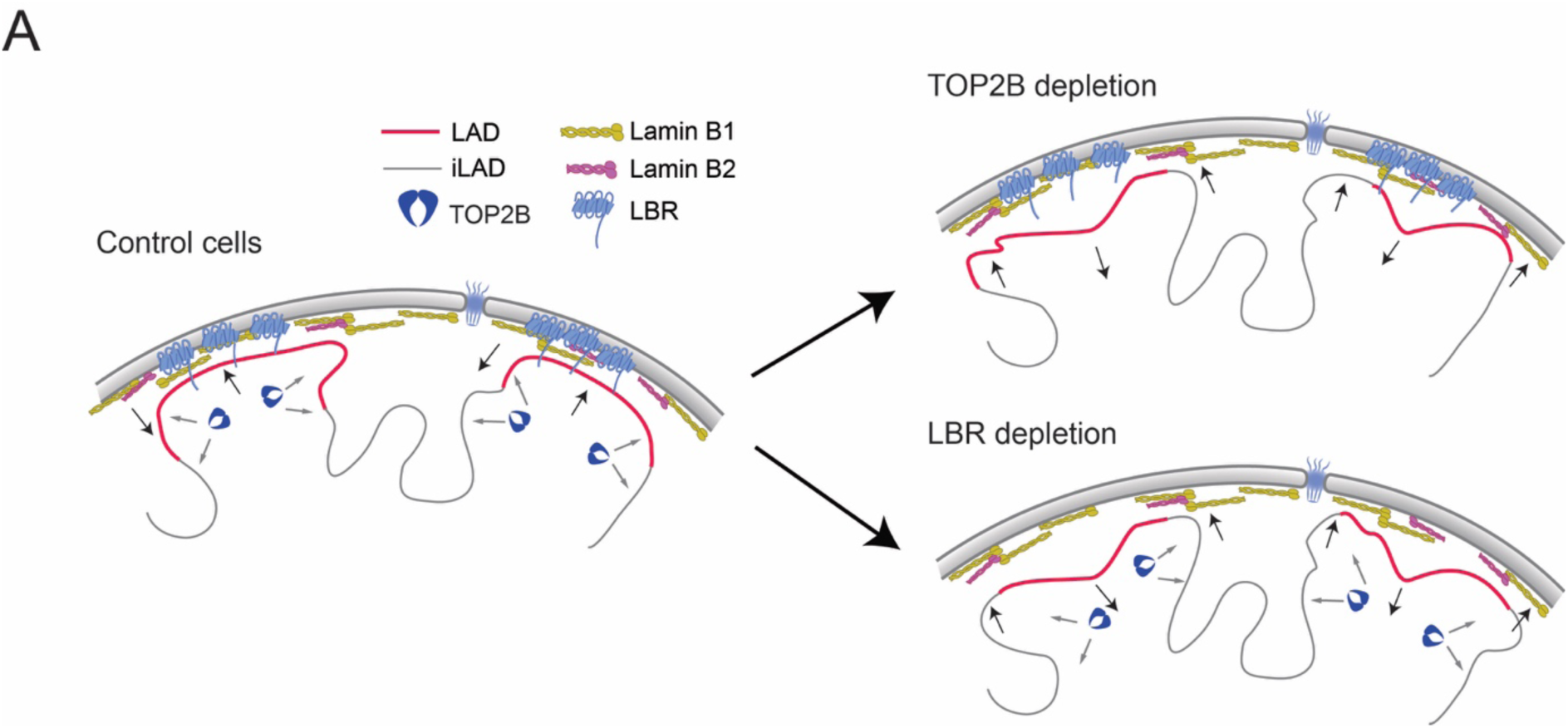
A) Possible model on how TOP2B and LBR can reinforce LAD/iLAD partitioning: LBR tethers LADs at the nuclear periphery while TOP2B maintain iLAD away from the NL. For all correlation scatter plots: the blue line represents a linear model; the black line represents the diagonal. Pearson correlation and p-value are shown in the plots.

One possible mechanistic speculation is that a low abundant, fast-acting subpopulation of TOP2B present in LADs (**Figure 3B**) helps to maintain a topological state optimal for LBR binding. Since topoisomerase cannot operate on constrained DNA (32,33), such a model would imply that LBR does not constantly constrain chromatin, but TOP2B could control its transient interactions and operate on unconstrained chromatin. This model would be consistent with data showing that genome-NL interactions are highly dynamic even in a single interphase (73).

In a second model, TOP2B helps LBR to bind chromatin at the mitotic exit, and the two proteins then segregate into two distinct genome compartments in interphase. A recent paper has shown that TOP2 helps to maintain an untangled and compartmentalised genome after mitotic exit (45). LBR is one of the first proteins to leave chromatin prior to NE disassembly and mitotic entry (74). It therefore needs to re-bind chromatin as cells exit mitosis and chromosomes begin to decondense. TOP2B activity may be required for optimal tethering at mitotic exit. In this model, the decatenation and relaxation activity of TOP2B might be necessary.

These models could also explain the topological preferences of LBR in DNA binding described *in vitro* (64). Further experiments will be needed to understand whether any of these models is a possible mechanism.

Despite the high similarity in the patterns of gain and loss of genome-NL interaction following loss of LBR or TOP2B, it is undeniable that depletion of the tether has a substantially more potent effect than the loss of topoisomerase. We have several explanations for this observation. First, LBR may play a critical role that cannot be replaced by any other tether in the nucleus. Differently, the depletion of TOP2B might be compensated by the activity of other topoisomerases like TOP2A or TOP1. Second, LBR has a nucleoplasmic portion with multiple domains dedicated to interactions with different chromatin components and features like histone modifications, DNA, lamins and HP1 (62,63). Thus, TOP2B might regulate only a specific aspect of the complex binding properties of LBR, and thus, the effect of TOP2B loss is limited.

Finally, although several data, such as 1) the sensitivity of genome-NL interactions to TOP2B depletion, 2) the differential topological state between LAD and iLAD, and 3) the DNA topology-dependent binding properties of LBR in vitro, suggest a potential role for DNA topology in controlling genome-NL interactions, the direct role of this biophysical property of chromatin in the processes described above remains to be demonstrated. Our data suggest that simple chromatin relaxation and dynamic supercoil removal are not sufficient to promote changes in genome-NL interactions. It is possible that DNA supercoils restrained by protein-DNA interactions in specific cell cycle stages, such as the ones typical of NE tethering, might be the key players in this phenomenon. In addition, the ability of topoisomerase II to entangle or disentangle the genome needs to be considered. Further studies will be required to understand which peculiar function of TOP2B is required for proper genome partitioning between the NL and the nuclear interior.

## SUPPLEMENTARY FIGURE LEGENDS

**Supplementary Figure 1.**
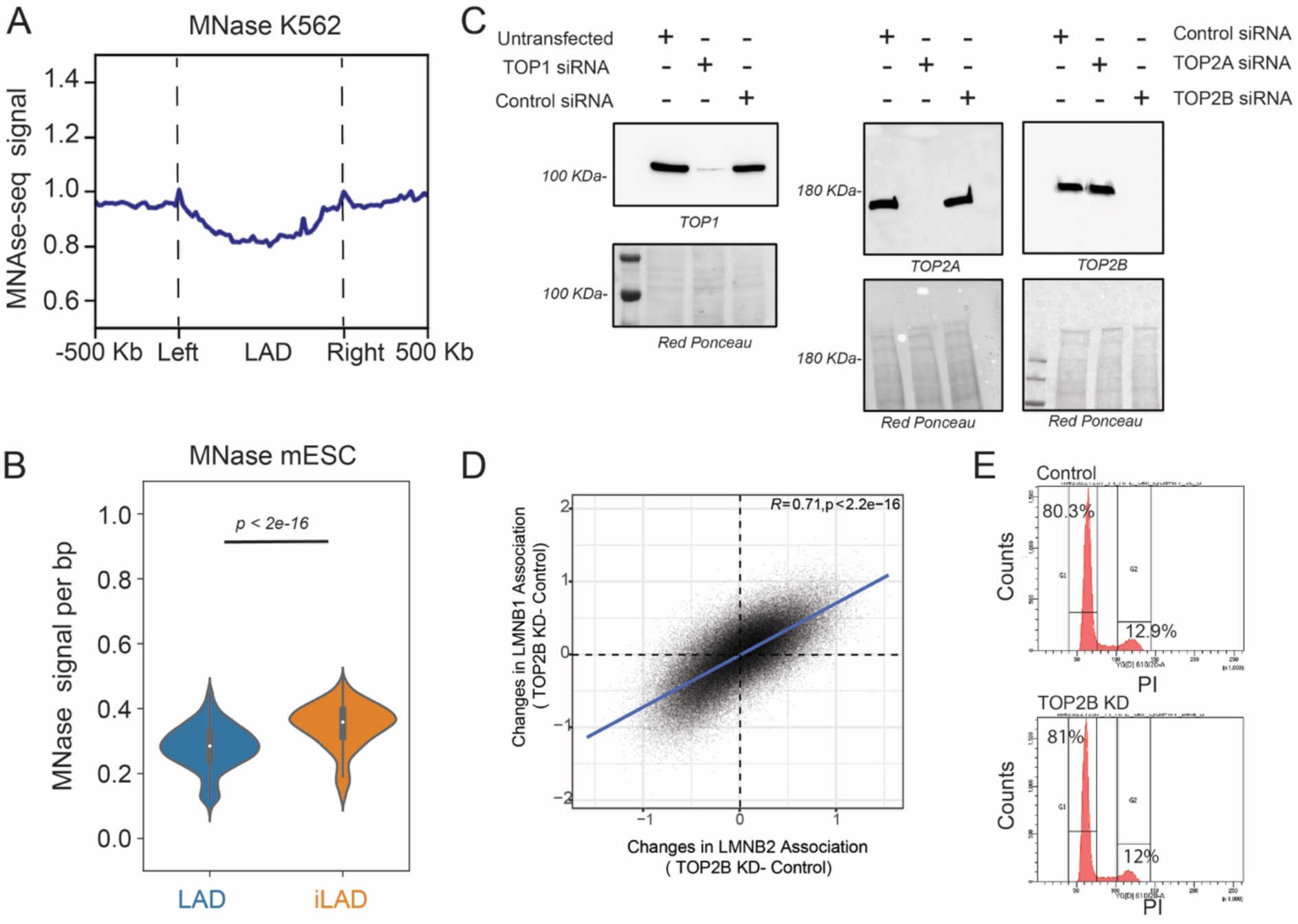
Mnase signal in LAD and iLADs and validation of topoisomerases knockdowns. A) Metagenic profile at LADs for MNAse signal in K562. LADs were scaled at the same length and the 500 kb upstream and downstream of LAD borders are shown. Data are from ENCODE (57). B) MNASE signal per bp for LAD and iLAD in mESC. P values are according to Welch’s T-test. Outlier values from top 0.5 and bottom 0.5 percentiles were excluded from plotting. Data are from (3). C) Western blot analysis to measure protein levels for TOP1, TOP2A, and TOP2B following siRNA-mediated depletions. Red Ponceau staining is used as a loading control. D) Correlation scatter plot of 20-kb genomic bins for differential LMNB2 and LMNB1 scores for TOP2B depletion (TOP2B knockdown - control). Results are from two biological replicates. The blue line represents a linear model. Pearson correlation and *p*-value are shown in the plot. E) Cell cycle analysis for Control and TOP2B depleted cells, by PI staining.

**Supplementary Figure 2.**
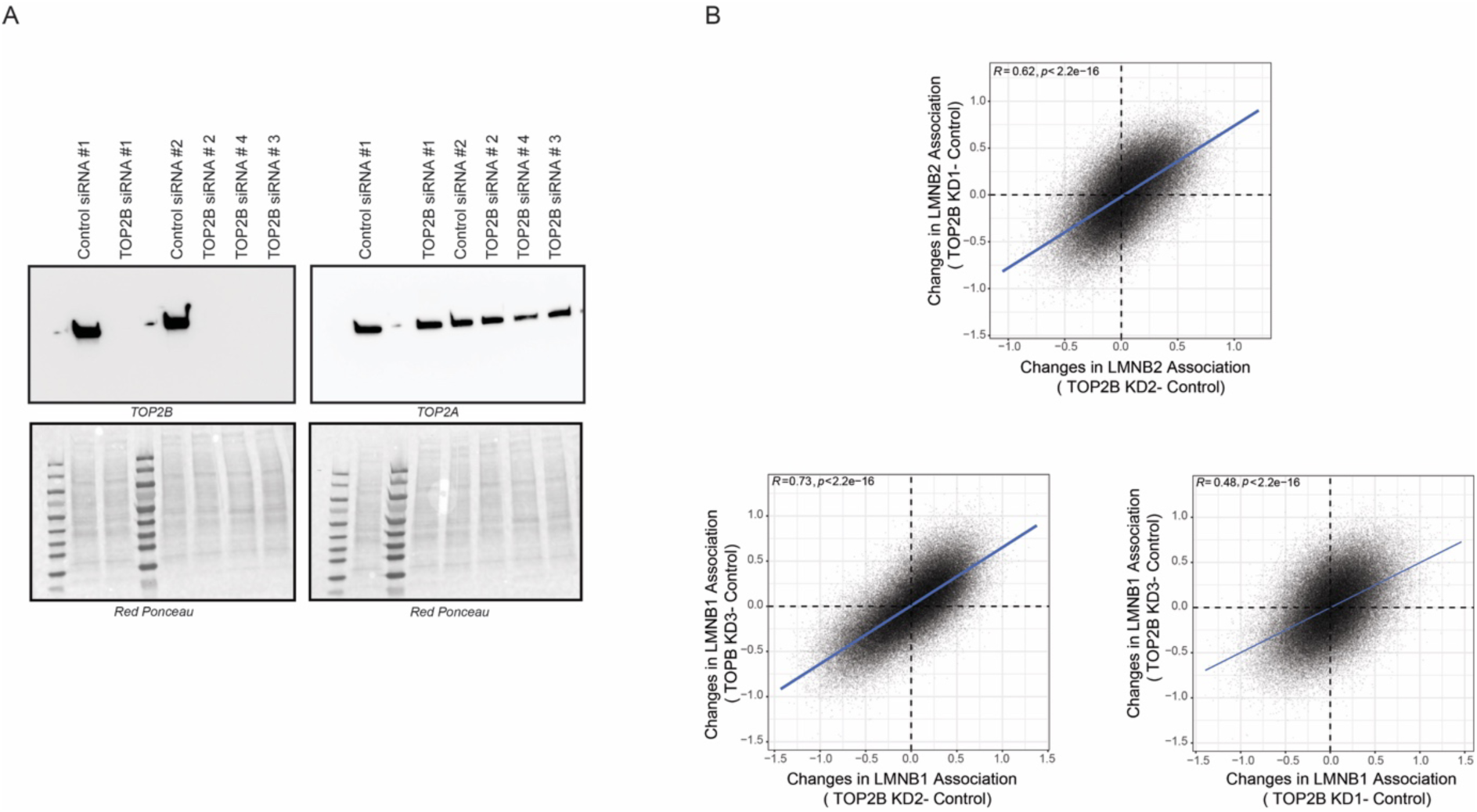
The effects of different TOP2B siRNAs show high correlations. A) Western blot analysis to detect TOP2B and TOP2A levels using different TOP2B siRNAs. Red ponceau was used as loading control. siRNA #4 was not used for further studies as it partially targets TOP2A. B) Correlation scatter plot of 20 kb genomic bins for differential LMNB2 and LMNB1 scores for TOP2B depletion (TOP2B knockdown - control) for three different siRNAs. Results are from two biological replicates. The blue line represents a linear model. Pearson correlation and *p*-value are shown in the plot.

**Supplementary Figure 3.**
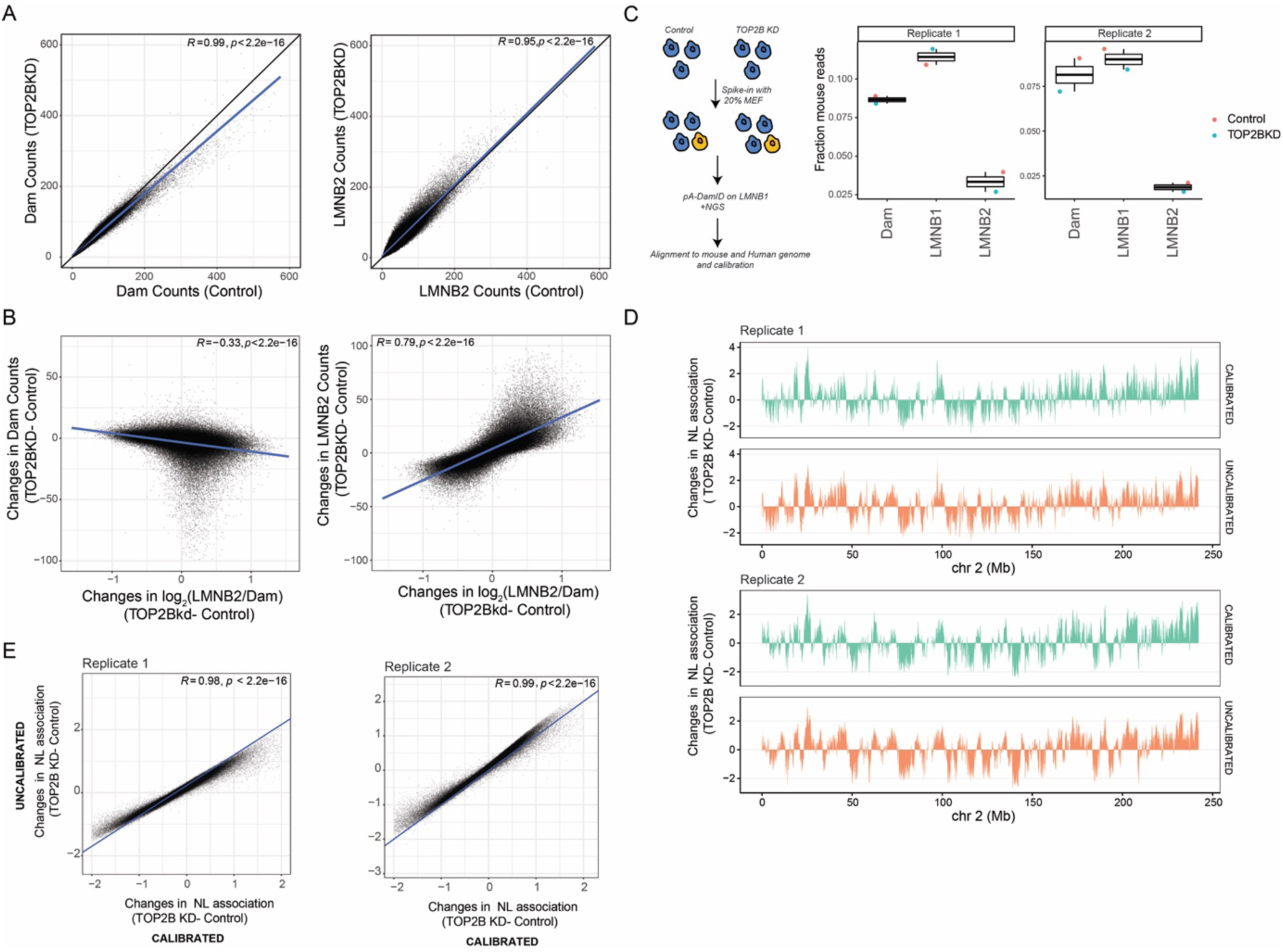
Controls of pA-DamID procedure. A) Correlation scatter plot for the “Dam only” signal (left panel) and “LMNB2 only” signal (right panel) for Control and TOP2B depleted cells. B) Correlation scatter plot for differential normalized log_2_ LMNB2/Dam score and differential Dam “only” or LMNB2 “only” for Control and TOP2B depleted cells. Data are from three biological replicates. C) *Left:* scheme explaining how calibration with mouse reads was performed. *Right:* Fraction of mouse reads in each sample (Dam, LMNB1, LMNB2) for each replicate. Note: LMNB2 antibody poorly performed in mouse cells and cannot be used for calibrated chromatin-NL contact maps. D) Differential LMNB2 score tracks (TOP2B depletion-control) for calibrated and uncalibrated data for each replicate. E) Genomewide correlation of Differential LMNB2 score (TOP2B depletion-control) for calibrated and uncalibrated data for each replicate. For all correlation scatter plots: the blue line represents a linear model; the black line represents the diagonal; Pearson correlation and *p*-value are shown in the plots.

**Supplementary Figure 4.**
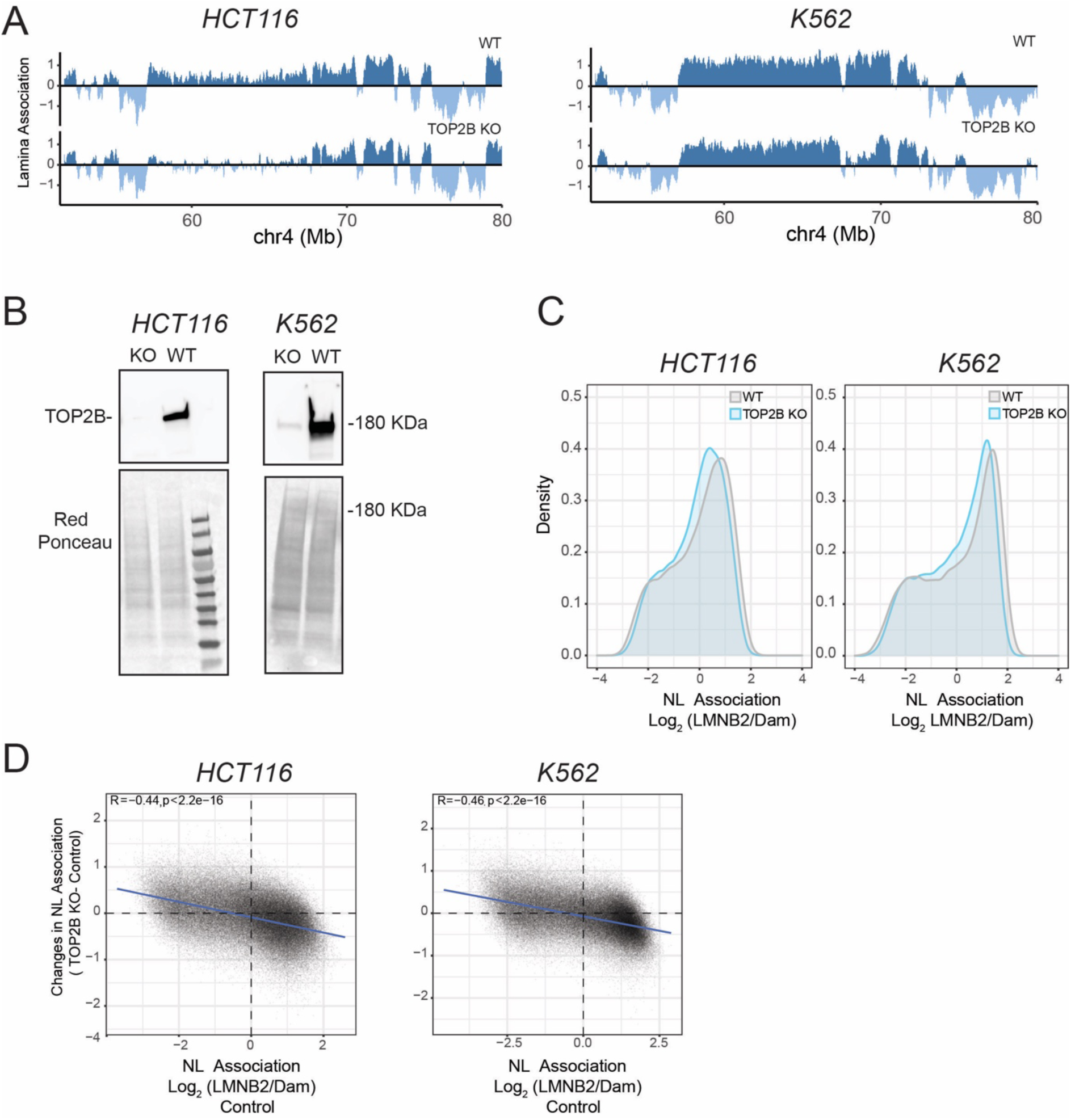
TOP2B control over genome-NL contact in additional cell lines. A) Representative genomic tracks of LMNB2 pA-DamID for control and TOP2B knockout in HCT116 (*left*) and K5632 (*right*). 20-kb bins were used. *B)* Western blot showing levels of TOP2B for the four different cell lines. Red ponceau staining was used as a loading control. C) LMNB2 signal distribution for control, and TOP2B knockout HCT116 cells (*left*) and K562 (*right*). D) correlation scatters plot of 20 kb genomic bins for LMNB2 score in control cells (x-axis) and differential LMNB2 score (TOP2BKO - control, y-axis) in HCT116 (*left*) and K562 (*right*). For all correlation scatter plots: the blue line represents a linear model. Pearson correlation and *p*-value are shown in the plots. The results are average of three biological replicates for HCT116 and two biological replicates for K562

**Supplementary Figure 5.**
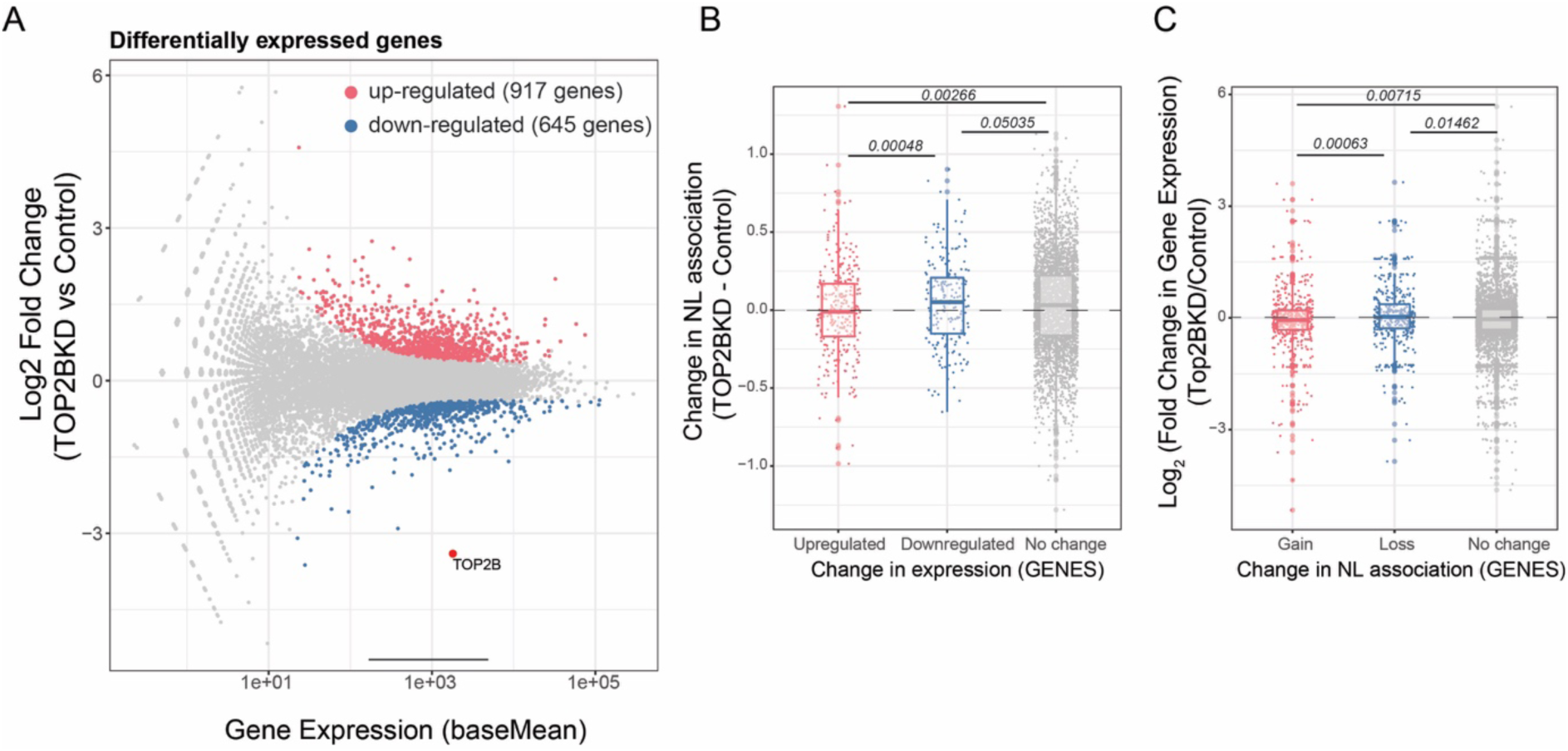
Effects of TOP2B depletion on gene expression. A) MA plot showing differentially expressed genes after TOP2B knockdown. Significant (*p adj* < 0.05) up and down-regulated genes are shown in red and blue respectively. TOP2B gene is highlited. B) Boxplot of differences in LMNB2 interaction score for up-regulated (red), down-regulated(blue), and unchanged genes (grey). C) Boxplot of log_2_(Fold Change) in gene expression for differentially attached (red), detached (blue) or stable genes (grey) following TOP2B depletion. To call differentially attached and detached genes a cut-off of +/− 0.4 in differential LMNB2 pA-DamID score was applied. P values are according to Wilcoxon’s test. Results are from two independent biological replicates.

**Supplementary Figure 6.**
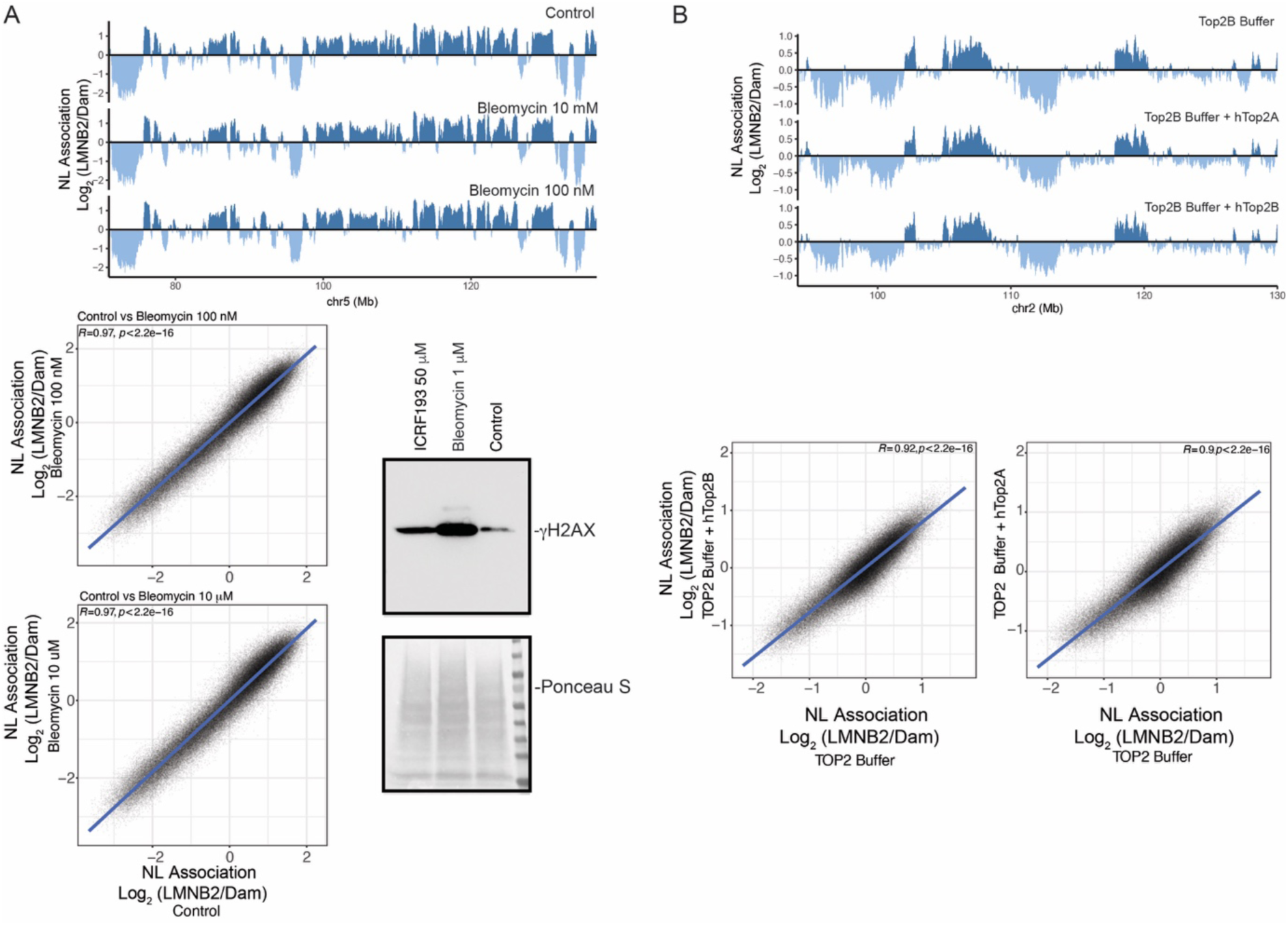
Dissipation of dynamic supercoils does not alter genome-NL interactions. A) *Upper part:* representative genomic tracks of LMNB2 pA-DamID for control RPE1 cells, and cells treated with 100nM and 10 μM Bleomycin for 3 hours. 20-kb bins were used. The antibody signal is normalized over a Dam-only control. *Bottom left* Correlation for LMNB2 signal between Control and bleomycin treatments. *Bottom right*: Western blot showing levels of γH2AX following three hours of treatment with bleomycin or ICRF193 (TOP2 catalytic inhibitors). Red ponceau staining was used as a loading control. B) *Upper part:* Representative genomic track of LMNB2 pA-DamID for control RPE1 permeabilized and resuspended in TOP2 Buffer or TOP2 Buffer plus human purified TOP2A or TOP2B. 20-kb bins were used for the analysis. The antibody signal is normalized over a Dam-only control. *Bottom part:* Correlation for LMNB2 signal between TOP2 Buffer and TOP2 Buffer plus topoisomerase enzyme.

**Supplementary Figure 7.**
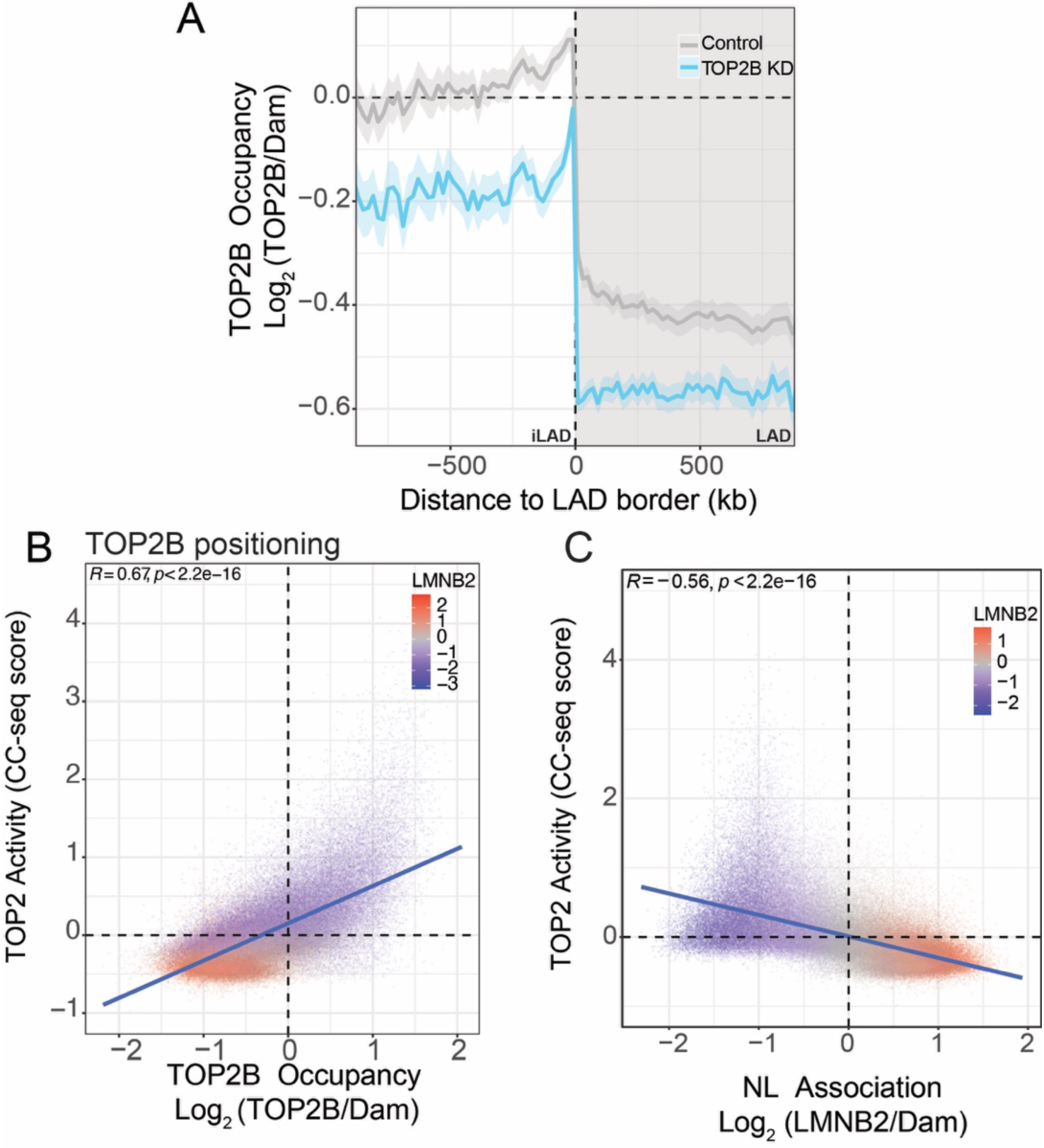
Genome wide-mapping of TOP2B positioning and activity. A) Average pA-DamID scores for TOP2B around LAD borders for RPE1 cells control (grey) and depleted for TOP2B (blue). The solid line and the shaded area represent the mean signal and 95% confidence interval of the mean, respectively. B) Correlation scatter plot between mapping data for TOP2B generated by pA-DamID and CC-seq signal that measure catalytically active TOP2. C). Correlation scatter plot between CC-seq signal that measure catalytically active TOP2 and lamina association measured by LMNB2-pA-DAmID. For all correlation scatter plots: the blue line represents a linear model; the black line represents the diagonal; Pearson correlation and *p*-value are shown in the plot. Results are from three biological replicates. CC-seq data are from (1).

**Supplementary Figure 8.**
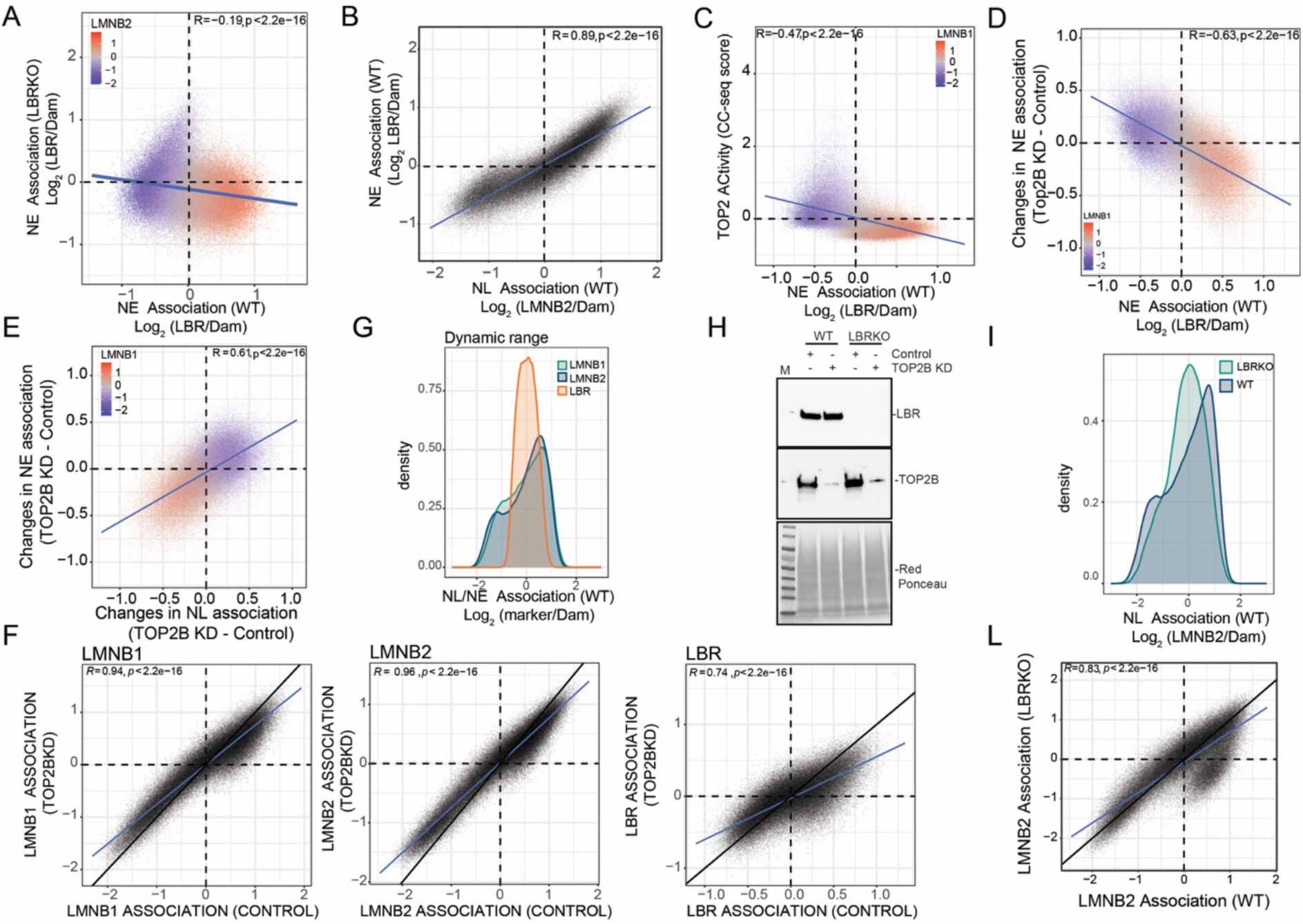
Mapping of genome-LBR interactions in control, TOP2B-depleted and LBRKO cells. A) Correlation scatter plot of 20 kb genomic bins for LBR signal between Control and LBR knockout RPE1 cells. B) Correlation scatter plot of 20 kb genomic bins for pA-DamID score for LMNB2 and LBR mapping in WT RPE1 cells. C) Correlation scatter plot of 20 kb genomic bins between LBR association and TOP2 activity in RPE1 cells. D) Correlation scatters plot of 20 kb genomic bins for LBR association score (x-axis) and differential LBR score (y-axis) for TOP2B depletion (TOP2BKD - control). E) Correlation scatter plot of 20 kb genomic bins for differential LMNB2 and LBR scores (TOP2B knockdown - control). F) Correlation scatter plot of 20 kb genomic bins for LMNB1, LMNB2 and LBR scores (Control vs TOP2B knockdown). G) Density plot showing dynamic ranges for LMNB1, LMNB2 and LBR mapping in RPE1 cells. H) Wester n blot showing the level of LBR and TOP2B in Control, TOP2B KD, LBR knockout, and co-depletion TOP2B and LBR. Red ponceau is used as loading control. I) Density plot showing LMNB2 signal distribution in WT and LBR knockout cells. L) Correlation scatter plot of 20 kb genomic bins for pA-DamID score for LMNB2 between Control and LBR knockout RPE1 cells. For all correlation scatter plots: the blue line represents a linear model; the black line represents the diagonal; Pearson correlation and *p*-value are shown in the plot. For A-I and L, results are from at least three biological replicates. For G, results are from two biological replicates and 4 technical replicates generated using two negative control siRNAs and two TOP2B-specific siRNAs.

**Supplementary Figure 9.**
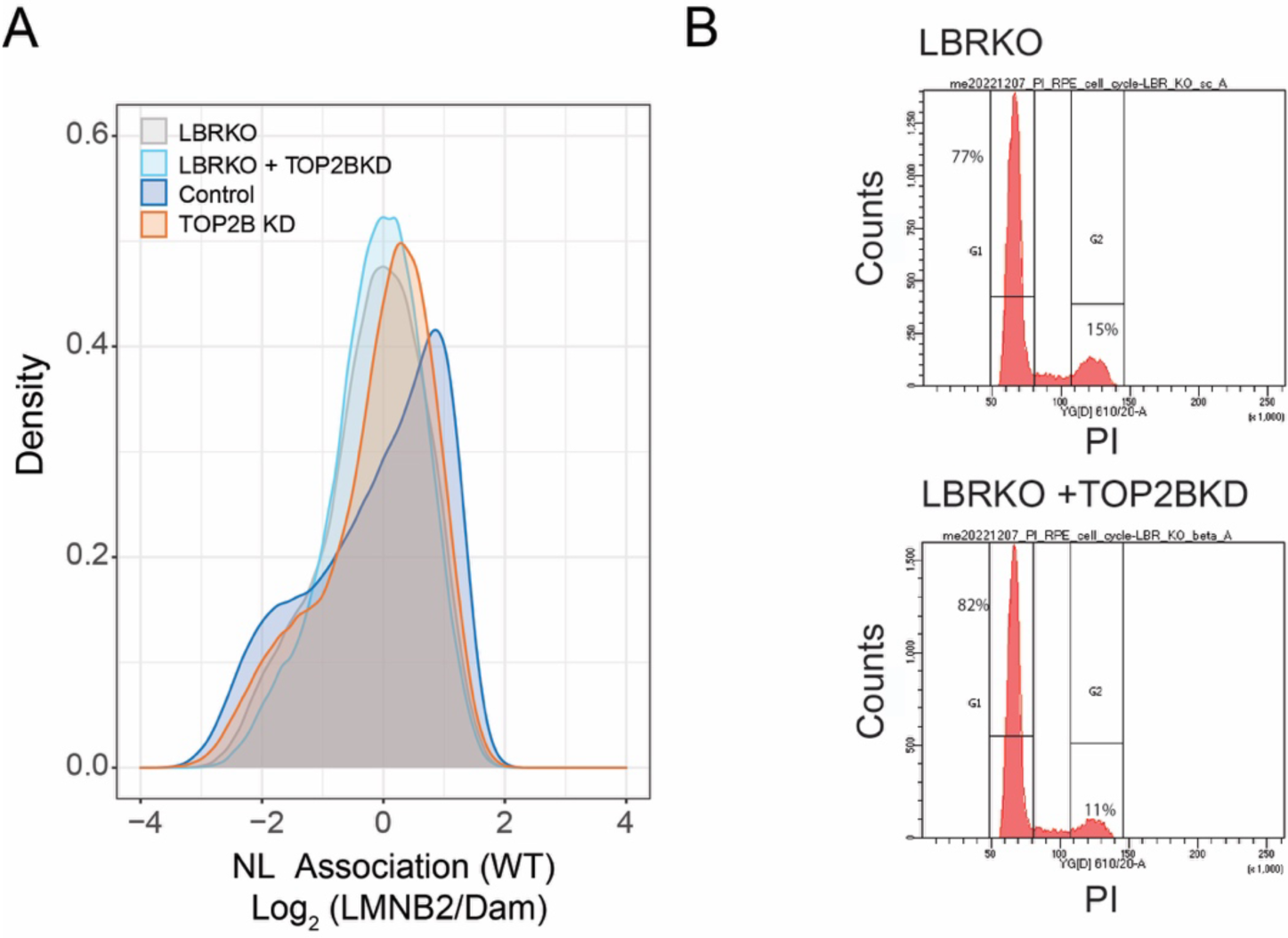
Co-depletion of TOP2B and LBR. A) Density plot showing LMNB2 signal distribution in WT, TOP2B depleted, LBR knockout and co-depleted (TOP2B knockdown + LBR knockout) RPE1 cells. B) Cell cycle analysis in LBR knockout cells with and without TOP2B, by PI staining.

**Supplementary Figure 10.**
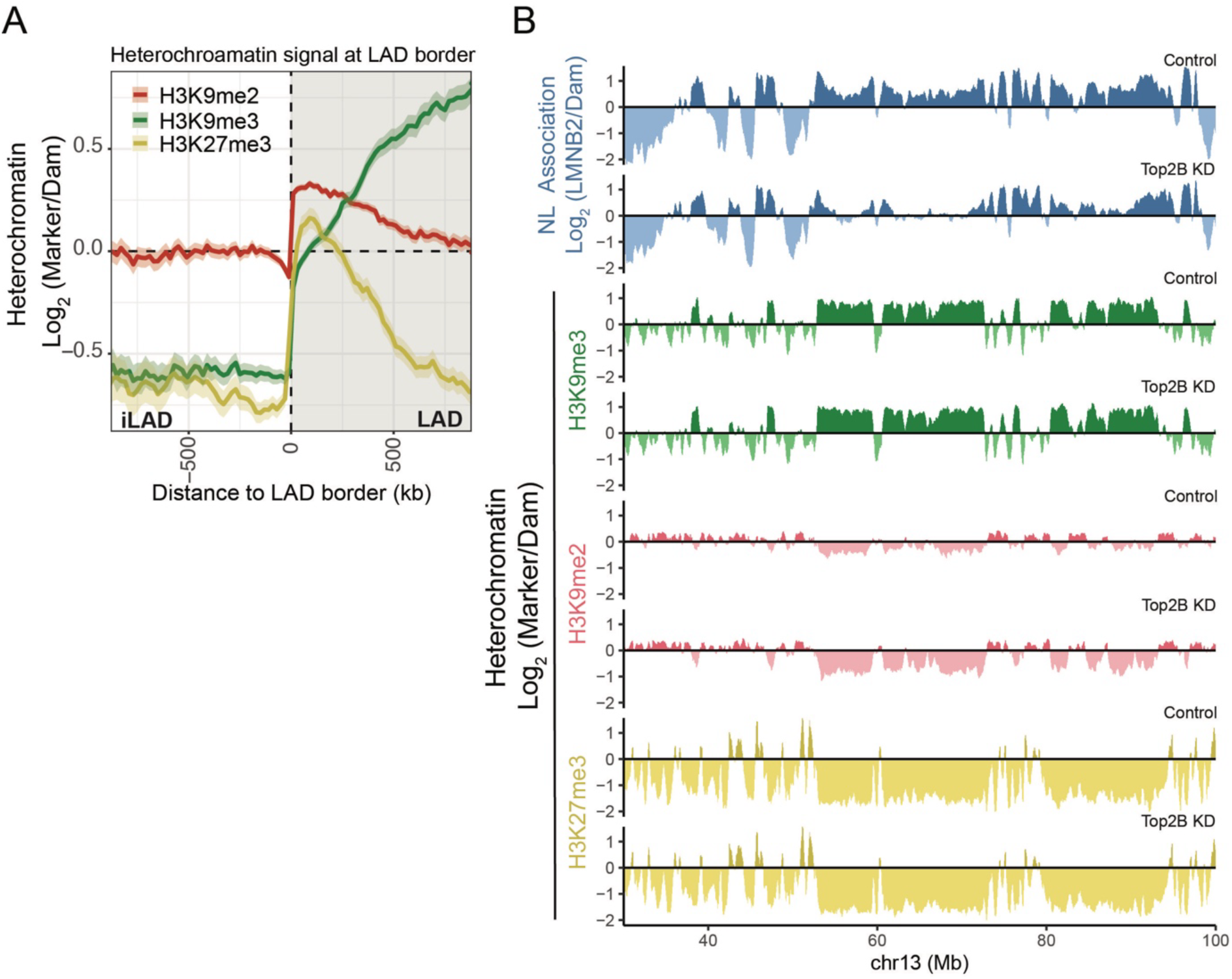
A) Average H3K9me3, H3K9me2 and H3K27me3 pA-DamID scores around LAD borders for WT RPE1 cells. The solid line and the shaded area represent the mean signal and 95% confidence interval of the mean, respectively B) Representative genomic track for pA-DamID signal for LMNB2 (blue), H3K9me3 (green), H3K9me2 (red), and H3K27me3 (yellow) for Control and TOP2B depleted cells. 20KB bins were used. The antibody signal is normalized over a Dam-only control.

## ACKNOWLEDGMENTS

We thank the NKI Genomics, Research High-Performance Computing and the Robotics & Screening core facilities for technical support; and members of our laboratories for stimulating discussions; Federico Comoglio for sharing R scripts. We thank Lise Dauban for critical reading of the manuscript, and members of our laboratory for helpful discussions. This work was supported by the European Union (ERC, GoCADiSC, 694466 to B.v.S.; AIRC-MSCA iCARE2 fellowship, 800924, MSCA-IF 838555, and Next generation EU-MUR MSCA Young Researcher to S.G.M; ERC, FuncDis3D, 865459 to E.d.W). R.M. is supported by by NWO Zwaartekracht (58588); N.G was supported by a UK Medical Research Council senior non-clinical fellowship (MR/J00913X/1) and funding from the MRC (MC_UU_00007/13). Views and opinions expressed are those of the authors only and do not necessarily reflect those of the European Union or the European Research Council. Neither the European Union nor the granting authority can be held responsible for them. Research at the Netherlands Cancer Institute is supported by an institutional grant of the Dutch Cancer Society and of the Dutch Ministry of Health, Welfare and Sport. The Oncode Institute is partially funded by the Dutch Cancer Society.

## AUTHOR CONTRIBUTIONS

S.G.M, B.v.S designed research. S.G.M, M.d.H, A.G.M, AJB, C.N, performed experiments. S.G.M, M.M, J.B, T.v.S performed bioinformatics analysis and analyzed data. S.G.M and B.v.S wrote the manuscript with input from all authors. S.G.M, B.v.S, E.d.W, R.M, N.G. acquired funding. All authors have read and approved the final manuscript.

## DATA AVAILABILITY

RNAseq and pA-DamID data generated in this work are available on GEO (accession code GSE277503, https://www.ncbi.nlm.nih.gov/geo/query/acc.cgi?acc=GSE277503). Labnotes and R scripts regarding this study can be found at https://osf.io/sp9ke.

## MATERIALS AND METHODS

### Cell lines and cell culturing

Two different hTERT immortalized RPE1 cell lines were used in this study. An RPE1 cell line that expresses the SunTag system (21,75) was used to perform TOP1, TOP2A and TOP2B depletions and the mapping of heterochromatin components. This cell line was mantained in DMEM (Gibco), FBS 10% and penicillin-streptomycin 1%. A second RPE1 cell line expressing and inducible Cas9 (76) was used for the generation of LBR knockout and all the other experiments. This cell line was mantained in DMEM:F12 (Gibco), FBS 10% and penicillin-streptomycin 1%. HCT116 were mantained in McCoy medium (Gibco) FBS10% and penicillin-streptomycin 1%. K562 cell lines were mantained in IMDM medium (Gibco), FBS10% and penicillin-streptomycin 1%. HCT116 and K562 TOP2B knockout cell lines were a kind gift from Vassilis Roukos. All cells were routinely checked for Mycoplasma contamination.

### Antibodies used

We used the following antibodies: LMNB2 (Abcam, ab8983), LMNB1 (Abcam, ab16048), LBR (Abcam, ab32535), H3K9me3 (Diagenode, C15410193), H3K9me2 (Active Motif, 39239), H3K27me3 (Diagenode, C15410195), TOP2B (Novus Biological, NB100-40842), TOP1 (Abcam, ab109374), TOP2A (Abcam, ab52934).

### Topoisomerases depletion

5,4 X10^5^ cells were transfected in suspension in a 10 cm-dish with 10 nM of siRNA and 1:1000 of Lipofectamin RNAimax reagent (ThermoFisher,13778075). 48 hours after transfection cells were detached and expanded in four 10-cm dishes and transfected again in the same conditions. All experiments were performed at 72 hours following the second round of transfection. Knockdown efficiency was routinely checked by western blot analysis. Two negative control siRNA were used: ON-TARGET plus Non-targeting Control Pool (Horizon Discovery, D-001810-10-05) and Silencer™ Select Negative Control No. 2 siRNA (ThermoFisher, 4390846). Cell lines transfected with these two siRNAs showed an high degree of correlation for chromatin-NL contacts (*R* >0.9), although some differences in LAD strenght could be detected. When possible, and as indicated in figure legends, the two datasets were averaged together. For TOP2B depletion three TOP2B-specific siRNAs were used: Dharmacon on target plus smart pool Human TOP2B siRNA (Horizon Discovery, L-004240-00-0005), Silencer Select TOP2B siRNA s106, (Thermofisher, 4390824) and Silencer Select TOP2B siRNA s108,(Thermofisher, 4390824). When possible, and as indicated in figure legends, the datasets from the siRNA pool and s108 were averaged together. For TOP1 depletion we used Silencer Select TOP1 siRNA (Thermofisher, S14304) and identical protocol to TOP2B depletion. For TOP2A depletion, we used Dharmacon on target plus smart pool Human TOP2A siRNA (Horizon Discovery, L-004239-00-0005). For this specific topoisomerase we performed a single knockdown as this was sufficient to completely block cell growth.

### pA-DamID

pA-DamID maps were generated as previously described (60). Briefly, one million of RPE1 cells were collected by centrifugation (500 g, 3 min) and washed sequentially in ice-cold PBS and digitonin wash buffer (D-Wash) (20 mM HEPES-KOH pH 7.5, 150 mM NaCl, 0.5 mM spermidine, 0.02% digitonin, Complete Protease Inhibitor Cocktail). Cells were rotated for 2 hours at 4 °C in 200 μL D-Wash with 1:200 antibody dilutuon, followed by a wash step with D-Wash. When the antibody used was not produced in rabbit, cells were incubated with a solution of D-Wash buffer and Rabbit-anti-mouse IgG (1:200, Abcam, ab6709), followed by a wash step. This incubation was repeated with a 1:200 pA-Dam solution (equivalent to nearly 60 Dam units, determined by calibration against Dam enzyme from NEB, #M0222L), followed by two wash steps. Dam activity was induced by incubation for 30 min at 37 °C in 100 μL D-Wash supplemented with the methyl donor SAM (80 µM) while gently shaking (500 rpm). For every condition, another 1 million cells were processed in only D-Wash and, during Dam activation, incubated with 4 units of Dam enzyme (NEB, M0222L). We use Dam-only control samples to normalize for DNA accessibility and amplification biases as described (77). Genomic DNA was isolated and processed similarly to DamID, except that the DpnII digestion was omitted. 65-bp reads were sequenced on HiSeq 2500 or 100 bp reads were sequenced on Novaseq platform. For Hiseq libraries library preparation was performed as previously described (60). For Novaseq libraries we performed the following modified protocol: genomic DNA was isolated (Bioline, BIO-52067) and ∼ 500 ng were digested with DpnI (10 U, NEB, R0176L) in CutSmart Buffer 1X (8h 37°C, 20 min 80°C) in a total volume of 10 µL. A-tailing was performed by adding 5 µL of the A-tailing mix (0.5 µL of Cutsmart buffer 10X, 0.25 µL Klenow 50 U/µL (NEB, M0212M), 0.05 µL dATP 100 mM, 4.2 µL H2O) and incubation 30 min at 37°C followed by 20 min at 75°C. Adapters were ligated by adding 15 µL of the ligation mix (3 µL T4 Ligase Buffer 10X, 0.5 µL T4 ligase (5 U/µL, Roche, 10799009001), 0.25 µl of x-Gene Stubby Adapter 50 mM (IDT), 11.25 µl H2O) and incubating samples 16h at 16°C and 10 min at 65°C. Finally, the Methyl Indexed PCR was performed by mixing 4 µL of ligated DNA with x-Gen Dual combinatorial Indexes (IDT) (125 nM final concentration) and MyTaq RedMix (Bioline, BIO-25048) in a final volume of 40 µL. The following PCR program was used: 1 X (1 min 94°C), 14 X (30 sec 94°C, 30 sec 58°C, 30 sec 72°C), 1X (2 min 72°C).

To perform calibrated pA-DamID we spiked-in RPE1 cells with 20% mouse embryonic fibrobalsts before proceeding to pA-DamID. For this specific experiment we performed LMNB1 pA-DamID as this antibody performed well in mouse cells, differently from LMNB2 antibody (see **Supplementary Figures 3C**).

### Bioinformatic analysis of pA-DamID data

pA-DamID data was processed as described (78). Briefly, the adapter sequence was trimmed with cutadapt 1.11 before mapping the remaining genomic DNA sequence to hg38 with bwa mem 0.7.17. The following steps were performed with custom R scripts. Reads with a mapping quality of at least 10 and overlapping the ends of a GATC fragment were counted in 20-kb genomic bins. Counts were normalized to 1 million reads per sample and a log_2_-ratio over the Dam control sample was calculated with a pseudo count of 1. At least 2 biological replicates were generated for every experimental condition and the average score was used for downstream analyses.

#### Differential LAD calling

Differential LADs calling was performed as previously described (60). Briefly, to call differential LADs we converted the log_2_ (LMNB2 : Dam) data to z-scores (mean of zero and a standard deviation of one) to account for differences in dynamic range in pA-DamID signals between experiments, thus keeping the data distribution identical. LADs were determined in the control sample using a hidden Markov model on the average NL interaction profile between biological replicates (https://github.com/gui11aume/HMMt). The LAD score was defined as the mean signal of the z-scaled data tracks. Significant changes were called using a modified limma-voom approach (65). Significance was defined as a Benjamini–Hochberg adjusted P-value lower than 0.05.

#### Analysis at LAD borders

LADs as described in the section “differential LAD calling” were used to visualize the consensus NL interaction profile around LAD borders. To calculate average signal at the LAD border we used 20 kb binned genomic tracks from bTMP-seq (IP/Input) and pA-DamID (Log_2_(Ratio)). The flanks of the LADs were extracted from the LAD models and used as LAD borders. LAD flanks within 50 kb of the chromosome ends were excluded as these are not actual LAD borders but simply chromosome ends. Next, for every 20 kb genomic bin kb, the closest LAD border was determined. Relative positioning from the closest LAD border was used to calculate the mean score and the 95%-confidence interval of the mean. Taking the closest LAD border for every genomic bin ensures that each position is only considered once, even when a position is close to multiple LAD borders. For this analysis, LADs and iLADs smaller than 50 kb were excluded as these are only composed of two normalized bins (of 20 kb each). These small domains would therefore only provide a single data point at one side of the LAD border, and likely only increase the noisiness of the resulting consensus profile.

#### Calibration of pA-DamID data

Calibrated pA-DamID data was processed similar to uncalibrated data, except that the reads were aligned to the human (hg38) and mouse (mm10) genomes at the same time and that genomic bins of 250 kb were used instead of 20 kb. The latter was necessary because the number of mouse reads was limited and larger bins resulted in more stable scaling estimates. The filtering step based on a mapping quality of 10 ensured that only unique alignments to either the human or mouse genome were used in the downstream analyses. After determining the log_2_-ratio over the Dam control using the combined genome reference, scaling factors were calculated to convert the mouse log_2_(ratios) to a z-score (see the section “different LAD scaling”). All mouse spike-ins were from the same biological sample, and this scaling ensured that all mouse profiles were normalized to the same dynamic range. Next, for each sample, the same scaling factors were used to scale the human log_2_(ratios). In contrast to uncalibrated data, with this approach quantitative differences in dynamic range between experiments due to increased or decreased protein-DNA interactions could be detected.

### In vivo and ex vivo chromatin relaxation

Bleomycin (Fisher Scientiifc, B2434-20MG) was dissolved in water, aliquoted and stored at −20°. Treatments were performed at 10 µM and 100 nM concentrations for thee hours before processing cells for pA-DamID. For *ex vivo* experiments, cells were permeabilized according to pA-DamID protocol and then resuspended in Human topo II assay buffer (Inspiralis, HTA202) and complemented or not with 20U of Human Topo II alpha (Inspiralis, HT205) or 20U of Human Topo II beta (Inspiralis, HTB205). Enzymes activities were checked with a plasmid relaxation assay.

### bTMP-seq

bTMP-seq was performed as previously described (52) with some modifications necessary for next generation sequencing. Cells or control genomic DNA were treated with 500 μg/ml bTMP for 20 min at room temperature in the dark. bTMP was UV cross-linked to DNA at 365 nm UV 800mJ/cm2 (UVP’s CL-1000). DNA was purified from cells using SDS and proteinase K digestion followed by phenol-chloroform-isoamyl alcohol extraction. DNA was fragmented by sonication to a range of 200-500 bp fragments. The bTMP−DNA complex in TE was immunoprecipitated using avidin conjugated to magnetic beads (Dynabeads MyOne Streptavidin Invitrogen, 65001) overnight at 4 °C. Beads were washed sequentially for 5 min each at room temperature with TSE I (20 mM Tris, pH 8.1, 2 mM EDTA, 150 mM NaCl, 1% Triton X-100 and 0.1% SDS), TSE II (20 mM Tris, pH 8.1, 2 mM EDTA, 500 mM NaCl, 1% Triton X-100 and 0.1% SDS) and buffer III (10 mM Tris, pH 8.1, 0.25 M LiCl, 1 mM EDTA, 1% NP40 and 1% deoxycholate). Beads were then washed twice with TE buffer for 5 min. bTMP enriched DNA was then “on-bead” end-repaired and A-tailed prior to ligation of Illumina sequencing adapters. Adapter-ligated bTMP-bound DNA was then eluted from the Dynabeads in 25 μl water by heating for 10 min at 98°C. Library preparation was continued using the NEB NEBNext® UltraTM II DNA Library Prep Kit for Illumina (NEB #E7645S) starting at “Step 4 : PCR amplification of adapter-ligated DNA” where 10 μl of the DNA was used. Final libraries were sized and quality controlled on a D1000 Tapestation tape (Agilent). Single-end DNA-sequencing of 50 bp read length was performed on Illumina Hi-Seq 4000 (UMC, Amsterdam).

### RNA-seq

RNA was extracted using an RNAeasy mini kit from Qiagen (#74104). One million cells were harvested, washed once in cold PBS, resuspended in 600 µL of RLT lysis buffer, and stored at −80°C. Libraries were prepared using the TruSeq® RNA LT kit and TruSeq RNA Single Indexes (Illumina). We sequenced libraries with single-end 65-bp reads on a HiSeq 2500 platform. We sequenced approximately 30 million reads for every condition. Two independent biological replicates were generated. RNA-seq reads were subjected to quality control using FastQC v0.11.6. Reads were aligned to the human reference genome (GRCh38, GRCh38_no_alt_analysis_set_GCA_000001405.15; https://www.encodeproject.org/files/GRCh38_no_alt_analysis_set_GCA_000001405.15/) using STAR 2.5.4a (79) with parameters --clip5pNbases 0 -- outWigStrand Unstranded. Gene-level count tables were generated while mapping using Gencode v24 primary assembly annotations.

### Analysis of publicly available data

#### CC-seq

Raw CC-seq data for K562 cells were downloaded from GSE136943 (1). The data was realigned to the GRCh38 human reference genome using the terminalMapper v4.1 pipeline (https://github.com/Neale-Lab/terminalMapper) as described in the original study. The resulting base-pair resolution profiles for Watson and Crick strands were merged together, binned into 20 kb bins and normalised using the total number of reads. We used these 20 kb binned profiles to compute the difference between VP16 treated and control RPE1 cells and obtained an estimate of catalytically engaged TOP2.

#### Nucleosome mapping

Processed MNase and chemical nucleosome positioning data for mouse embryonic stem cells were downloaded from GSE82127 (3). Constitutive LAD and inter-LAD (cLAD and ciLAD, respectively) regions in the mouse cell types were obtained from GSE17051 (55). Nucleosome signals for MNase and chemical mapping were calculated for each cLAD and ciLAD region using the “intersect” and “groupby” functions from BEDTools v2.27.1 (80). Statistical significance of the difference between nucleosome signals in cLAD and ciLAD regions was assessed using a T-test. Outlier values from top 0.5 and bottom 0.5 percentiles were excluded from plotting. Linker DNA lengths in cLAD and ciLAD regions for chemical nucleosome mapping were calculated as described in the original study (3). First, the nucleosome positions located in the cLAD and ciLAD regions were identified using the “intersect” function from BEDTools. Second, the distance between two consecutive nucleosome centres was calculated. Lastly, in case the distance was larger than 100 bp, 147 bp was subtracted. Linker DNA lengths between 1 and 100 bp were used for plotting.

MNase-seq data for K562 cells was downloaded from GSM920557 (57) and lifted to the GRCh38 human reference genome using UCSC liftOver. Pile-up of the MNAse-seq data over K562 LADs (identified by pA-DamID) was calculated using the “computeMatrix” function from deepTools v3.4.2 (81) in scale-regions mode with parameters ‘-bs’ set to 10000, ‘--beforeRegionStartLength’ set to 500000, ‘--regionBodyLength’ set to 1000000, ‘--afterRegionStartLength’ set to 500000, ‘--missingDataAsZero’, and ‘--skipZeros’. Pile-ups were visualised using the “plotHeatmap” function from deepTools. *OIS data.* RNAseq counts for OIS experiments was downloaded from GSE130306 (2). Genome-NL interactions maps during OIS wad downloaded from GSE76605 (4) and re-aligned to GRCh38.

## Notes

### Competing Interest Statement

The authors have declared no competing interest.

